# Multivariate analysis of strawberry experiments: where are we now and where can we go?

**DOI:** 10.1101/2020.12.30.424876

**Authors:** Tiago Olivoto, Maria Inês Diel, Denise Schmidt, Alessandro Dal’Col Lúcio

## Abstract

The multi-trait genotype-ideotype distance index (MGIDI) was used to select superior treatments in experiments with strawberries. Twenty-three productive, qualitative, physiological, and phenological traits with negative and positive desired gains were accessed in 16 treatments, a combination of two cultivars (Albion-neutral days, and Camarosa-short days), two transplants origins (National and Imported), and four organic substrates mixes (Crushed sugarcane bagasse, burnt rice husk, organic substrate, and Carolina commercial substrate). Our results suggest that most of the strawberry traits are influenced by the cultivar, transplant origin, cultivation substrates, as well as by the interaction between cultivar and transplant origin. The MGIDI index indicated that the Albion cultivar originated from imported transplants grown in substrates where the main component (70%) is burnt rice husk provides desired values for 20 of a total of 22 traits, which represents a success rate of ~ 91% in selecting traits with desired values. The *strengths and weakness view* provided by the MGIDI index revealed that the looking for an ideal treatment should direct the efforts on improving the water efficiency use and reducing total acidy of fruits. On the other hand, the strengths of selected treatments are mainly related to productive precocity, total soluble solids, and flesh firmness. The MGIDI index provides a unique, robust, and easy-to-handle process, standing out as a powerful tool to develop better treatment recommendations for experiments with strawberries when multiple traits are assessed.

## 1. Introduction

Strawberry productivity has grown in recent years, as it is a crop with high economic benefits (Singhalage et al., 2019), in addition to the nutraceutical and sensory characteristics of its fruits (Akhatou and Fernández-Recamales, 2014). The world harvested area with strawberry ranged from 363.398 ha in 2016 to 372.361 ha in 2018. As a result, fruit production also increased, from 7.901 million metric tonnes in 2016 to 8.34 million metric tonnes in 2018, and in the last year, the average yield reached 182.3 Mg ha^−1^. The world’s largest strawberry producers are the United States and China (FAO, 2020).

Strawberry production and productivity are changed every year. New research seeks ideal growth conditions for cultivation, in addition to genotypes with high yield and fruit quality. Besides, the planting of high-quality seedlings is crucial to express the genetic potential in the cultivation environment (Park et al., 2020).

The cultivated strawberry has a limited genetic basis (Hardigan et al., 2018; Gaston et al., 2020) and because of this, it makes it difficult to quickly identify important target traits for the greatest success in improving the strawberry (Gaston et al., 2020). Generally, it is sought for cultivars that present good productivity, that is precocious, and with a high rate of production (Diel et al., 2020). Sensory characteristics such as total soluble solids content, total titratable acidity, and their relationship, are essential to winning over consumers, as well as physical coloring characteristics (Adak et al., 2017; Diel et al., 2018; Gaston et al., 2020). Besides, flesh firmness or resistance to penetration is essential to increase the shelf-life of fruits (Diel et al., 2018; Rahman et al., 2016).

In many cases, genotypes with optimal chemical characteristics, such as high flavor and aroma, maybe unproductive, due to the cultivation environment or genotype characteristics, and are not ideal from the point of view of the producer. The cultivar Camarosa, for example, showed higher production, but lower sugar content and higher acidity (Diel et al., 2018), it may not be attractive for fresh consumption, for example, and attributed to the processing industry.

The selection of strawberry hybrids for consumption in natura or processed through multivariate analyzes to obtain characters of interest was performed by Barth et al. (2020). The authors calculated the selection index based on different weights adopted for the fresh and processed market, with the evaluated characteristics showing high variability between hybrids, and the greatest selection gains were obtained for production characteristics. Besides, the different weight assignments have resulted in different classifications of hybrids for fresh and processed consumption.

In this context, Olivoto and Nardino (2020) have proposed a selection index for selecting genotypes and/or recommending treatments based on information of multiple traits. The multi-trait genotype-ideotype distance index (MGIDI) allows a more efficient and accurate treatment recommendation based on desired or undesired characteristics for the crop studied (Olivoto and Nardino, 2020). Thus, the objective of this study was to evaluate and select the combination of the best factor combinations (cultivar, transplant origin, and substrate mixture) in strawberry production where most of the characteristics show favorable gains using the multi-trait genotype-ideotype distance index.

## 2. Material and methods

### 2.1. Location and cultivation environment

The experiment was conducted at the Federal University of Santa Maria, Frederico Westphalen campus (27°23’S, 53°25’O, 493 masl). The climate is Cfa according to Köppen’s classification, where the three coldest months of the year have temperatures of −3 to 18° C, with an air average temperature in the warmest month greater than or equal to 22°C, and precipitation uniformly distributed during the year (Alvares et al., 2013).

An open, substrate cultivation system was carried out inside an experimental greenhouse (20-m length, 10-m width, and 3.5-m height). The strawberry transplants were transplanted into white, 150-μm thickness tubular plastic bags, kept on wooden benches 0.8 meters above the ground. Drip irrigation and fertigation were performed. The frequency of irrigation and the formulation of fertigation was carried out according to Gonçalves et al. (2016). The nutrient dose for fertigation is presented in Supplementary Table S1. The electrical conductivity (EC) of the nutrient solution was 1.8 mS cm^−^1. The irrigation frequency as well as the time of each irrigation pulse were adjusted based on the solution drained from the substrate, monitoring the EC of the nutrient solution.

### 2.2. Plant material and experimental design

The experiment was conducted in a randomized complete block design with four replications in a threeway factorial treatment structure with two cultivars (Albion-neutral days, and Camarosa-short days), two transplants origins (National and Imported), and four organic substrates mixes (Crushed sugarcane bagasse, burnt rice husk, organic substrate, and Carolina^®^ commercial substrate), totaling 16 treatments (Table 1). Each experimental unit was composed of 8 strawberry plants. The water retention curve for different substrates used in the experiment is shown in Fig. 1. Sugarcane bagasse-based substrates presented a more pronounced reduction in moisture with low tensions (1 Kpa), which can be explained by the higher aeration space for these substrates (Supplementary Table S2).

**Figure 1:**
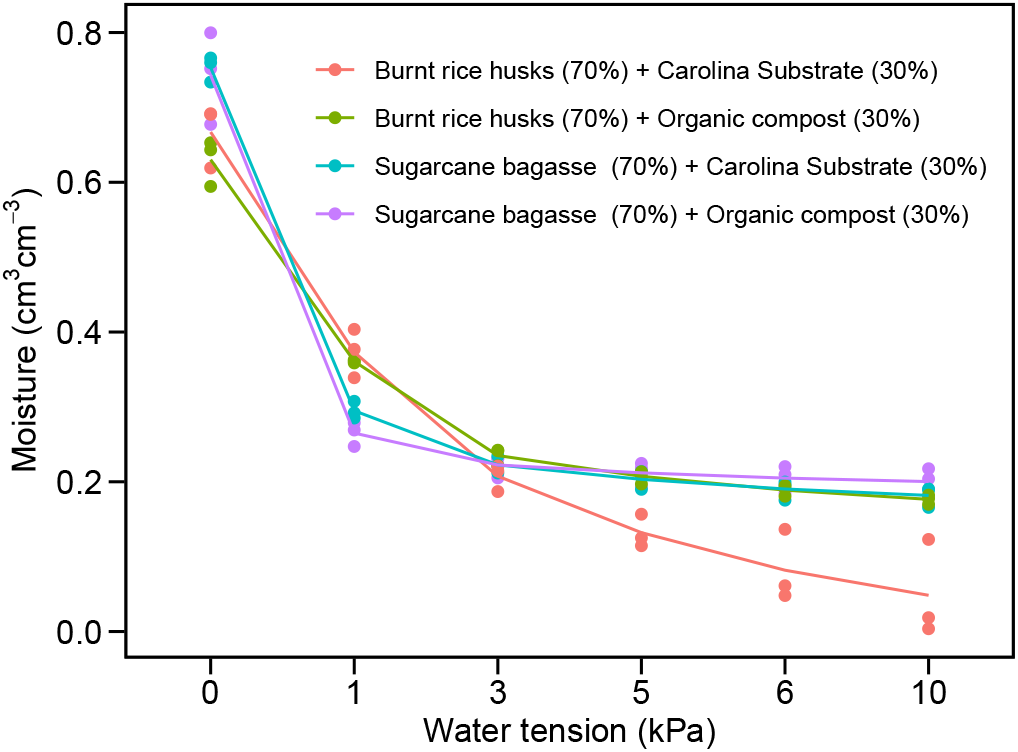
Water retention curve for different substrates used in the experiment.

**Table 1:** Treatment code used to characterize the experiment of two strawberry cultivars coming from two transplant origins grown cultivated on different substrates.

| Treatment | Cultivar | Origin | Substrate for cultivation |  |  |  |
| --- | --- | --- | --- | --- | --- | --- |
|  |  |  | Sugarcane bagasse (%) | Rice husk (%) | Organic (%) | Carolina<sup>®</sup> substrate (%) |
| T1 | Camarosa | National | 70 | 0 | 30 | 0 |
| T2 | Camarosa | Imported | 70 | 0 | 30 | 0 |
| T3 | Albion | National | 70 | 0 | 30 | 0 |
| T4 | Albion | Imported | 70 | 0 | 30 | 0 |
| T5 | Camarosa | National | 70 | 0 | 0 | 30 |
| T6 | Camarosa | Imported | 70 | 0 | 0 | 30 |
| T7 | Albion | National | 70 | 0 | 0 | 30 |
| T8 | Albion | Imported | 70 | 0 | 0 | 30 |
| T9 | Camarosa | National | 0 | 70 | 30 | 0 |
| T10 | Camarosa | Imported | 0 | 70 | 30 | 0 |
| T11 | Albion | National | 0 | 70 | 30 | 0 |
| T12 | Albion | Imported | 0 | 70 | 30 | 0 |
| T13 | Camarosa | National | 0 | 70 | 0 | 30 |
| T14 | Camarosa | Imported | 0 | 70 | 0 | 30 |
| T15 | Albion | National | 0 | 70 | 0 | 30 |
| T16 | Albion | Imported | 0 | 70 | 0 | 30 |

The transplants of National origin came from Agudo-RS (29°62’S, 53°22’O, 83 masl). The Imported transplants were produced in the Patagônia. Agrícola SA nursery, located in the municipality of El Maitén in Argentina (42°3’S, 71°10’O, 720 masl). The transplants of the cultivars Albion (National) and Camarosa (National and Imported) were transplanted on May 26, 2015. The cultivar Albion imported was transplanted on June 8, 2015.

### 2.3. Assessed traits

The harvests began in the maturation stage and were carried out twice a week. Through the production cycle, a total of 22 phenological, productive, physiological, and qualitative traits were assessed.

#### 2.3.1. Phenological traits

We evaluated the following phenological traits: phyllochron (PHYL, °C day leave^−1^ estimated as the inverse of the slope of the linear regression between the number of leaves in the crown against the accumulated thermal sum. A leaf was counted when the leaflets did not touch each other; number of days for the beginning of flowering (NDBF, days), computed when the first flower of the block was open; number of days for full flowering (NDFF, days), computed when all plants in the block had opened flowers; number of days for beginning of harvest. (NDBH, days);

#### 2.3.2. Productive traits

The fruits from each plot were classified as commercial and non-commercial and evaluated separately. We considered as non-commercial, fruits deformed or with weight less than 6 grams. In this stage, the following productive traits were assessed: number of commercial fruits (NCF, fruits plant^−1^); number of non-commercial fruits (NCF, fruits plant^−1^); total number of fruits (TNF, fruits plant^−1^); weight of commercial fruits (WCF, g plant^−1^); weight of non-commercial fruits (WNCF, g plant^−1^); total weight of fruits (TWF, g plant^−1^); average weight of commercial fruits (AWCF, g fruit^−1^); average weight of non-commercial fruits (AWNC g fruit^−1^); overall average weight of fruits (OAWF, g fruits^−1^); fruit yield (FY, Kg ha^−1^).

#### 2.3.3. Physiological traits

The water use efficiency (WUE, l^−1^) calculated the ratio between the amount of water used in the entire duration of the experiment and the total fruit production for each plant.

#### 2.3.4. Qualitative traits

Quality-related traits were assessed in three moments throughout the production cycle, to smooth possible punctual season effects. The average value of each plot was then used for the following traits: total titratable acidity (TA, mg citric acid 100 g^−1^) performed by titration with a standardized NaOH solution (0.1 mol L^−1^), total soluble solids (TSS, °Brix) using a manual refractometer (±2% accuracy), the ration between TSS and TA (TSS/AT) calculated using the quotient between the total soluble solids content and the titratable acidity; flesh firmness (FF), determined using a bench penetrometer with a 6 mm plunger.

Pulp coloration was evaluated by chroma, hue angle, and lightness in CIELCh color space (cylindrical coordinates) after conversion from the CIEL*a*b* color space (cartesian cordinates). First, fruit color was expressed as three values [L*, for the lightness from black (0) to white (100), a* from green (−) to red (+), and b* from blue (−) to yellow (+)]. L*, a*, and b* were determined with a colorimeter calibrated with a standard white ceramic plate. The conversion of a* and b* to C* (CHROMA, relative saturation) and h^°^ (H, angle of the hue in the CIELab color wheel) was done using the following formulas: 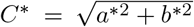; and *h*^°^ = arctan (*b*\*/*a*\*). The CIEL*a*b* lightness (L*) remained unchanged.

### 2.4. Defining an *ideal* treatment

In this crucial step, we define an *ideal* treatment, i.e., the one that would provide desired values for all studied traits. Based on previous knowledge, and assuming that a grower focuses on early production of strawberries, an ideal treatment should provide: (i) plants with high water use efficiency that presents a short period between planting and beginning of flowering/harvesting, with low phyllochron values, i.e., a high number of leaves per accumulated thermal sum (i.e., lower values for WUE, NDBF, NDFF, NDBH, and PHYL); (ii) high fruit yield with a higher number of commercial fruits with a high average weight of fruits (i.e., higher values for NCF, TNF, WCF, TWF, AWCF, OAWF, and FY); (iii) lower number of noncommercial fruits of low average weight (i.e., lower values for NNCF, WNCF, and AWNCF); and (iv) sweet, firm fruits with lower acidity and ideal external and internal red color (i.e., lower values for TA and higher values for TSS, TSS/TA, FIRM, L, CHROMA, and H).

### 2.5. Statistical analysis

#### 2.5.1. Estimated marginal means

Each trait was analyzed according to the following three-Way ANOVA cross-classification model.

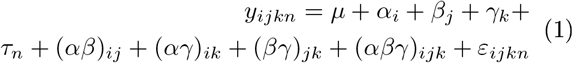

where *y_ijkn_* is the observed value in the *n*th block (*n* = 1, 2,…4) of the *i*th cultivar (*i* = 1, 2) from the *j*th origin (*J* = 1, 2) cultivated in the *k*th substrate (*k* = 1, 2,…, 4); *μ* is the the general mean; *α_i_* is the effect of the *i*th cultivar; *β_j_* is the effect of the *J*th origin; *γ_k_* is the effect of the *k*th substrate; *τ_n_* is the effect of the nth block; (*αβ*)_ij_ is the interaction effect between *α_i_* and *β_j_* (*αγ*)_*ik*_ is the interaction effect between *α_i_* and *γ_k_*; (*βγ*)_*jk*_ is the interaction effect between *β_j_* and *γ_k_*; (*αβγ*)_*ijk*_ is the interaction effect between *α_i_*, *β_j_* and *γ_k_*; and ϵ_*ijkn*_ is the error term associated with *y_ijkn_*. The error term *ε* is identically and normally distributed with zero mean and variance σ^2^, that is, *ε* ~ *N*(0, σ^2^).

A two-way table (X_*ij*_) containing the estimated mean for the three-way term in Eq. 1 in the rows for each trait in the columns was created. Here, *t* is the number of treatments *t* = 1, 2,…, *16* (2 × 2 × 4) and *ν* is the number of traits in the study *ν* = 1, 2,…, *23*.

#### 2.5.2. Reescaled means

Based on the knowledge of an *ideal* treatment, we reescaled X_*ij*_ to obtain *rX_ij_* as proposed by Olivoto et al. (2019).

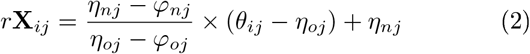

where *η_nj_* and *φ_nj_* are the new maximum and minimum values for the trait *j* after rescaling, respectively; *η_oj_* and *φ_oj_* are the original maximum and minimum values for the trait *j*, respectively, and *θ_ij_* is the original value for the *j*th trait of the *i*th treatment. For NNCF, WNCF, AWNCF, WUE, NDBF, NDFF, NDBH, PHYL, and TA in which lower values are desired, *η_nj_* = 0 and *φ_nj_* = 100 were used. For all the other traits in which higher values are desired, *η_nj_* = 100 and *φ_n_j* = 0 were used. In the rescaled two-way table (*rX_ij_*), all columns have a 0-100 range in which 0 is the most undesired value and 100 is the most desired value. Thus, the ideal treatment would be that with 100 for all traits after rescaling.

#### 2.5.3. Selecting the best treatment combinations

The MGIDI index (Olivoto and Nardino, 2020) was used to rank the treatments based on the desired values of studied trait. First, a factor analysis was computed with (*rX_ij_*) to account for the correlation structure and dimensionality reduction of the data. Eigenvalues greater than one were retained. Then, an Euclidean distance between the scores of treatments and the ideal treatment was computed as follows (Olivoto and Nardino, 2020):

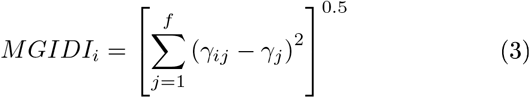

where *MGIDI_i_* is the multi-trait genotype-ideotype distance index for the *i*th treatment; *γ_ij_* is the score of the *i*th treatment in the *J*th factor (*i* = 1, 2,…,*t*; *j* = 1,2,…,*f*), being *t* and *f* the number of treatments and factors, respectively; and *γ_j_* is the *j*th score of the ideal treatment. The treatment with the lowest MGIDI is then closer to the ideal tratament and therefore presents desired values for all the *ν* traits.

The selection differential for all traits was computed considering a selection intensity of ~ 15%, i.e., the first two treatments with the lowest MGIDI index were selected.

The proportion of the MGIDI index of the *i*th treatment explained by the jth factor (*ω_ij_*) was used to show the strengths and weaknesses of the treatments and was computed as:

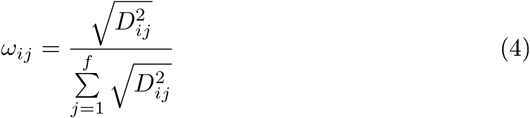

where *D_ij_* is the distance between the *i*th treatment and ideal treatment for the *j*th factor. Low contributions of a factor indicate that the traits within such a factor are close to the ideal treatment. Data manipulation and the index computation were performed in the R Software version 4.0.2 (R Core Team, 2020) using the package metan (Olivoto and Lúcio, 2020).

## 3. Results

### 3.1. Analysis of variance

The analysis of variance revealed significant (*p* ≤ 0.05) the main effect of substrate (SUB), origin (ORI), and cultivar (CUL) for 13, 10, and 18 traits, respectively (Supplementary Table S3). The two-way interaction term ORI-CUL was significant for 12 traits, while the interaction terms SUB-CUL and SUB-ORI were significant for the traits PF and L, respectively. The three-way interaction term (CUL-ORI-SUB) was only significant for TA (Supplementary Table S3).

### 3.2. Linear correlation, Loadings, and Factor delineation

The traits WCF. TNF, NCF, TM, TWF where positive and highly correlated (*r* > 0.75, *p* < 0.001) with each other (Supplementary Fig. S2). The negative correlations between these traits with WUE (—0.82 ≤ *r* ≤ −0.7,*p* < 0.001 −0.35 ≤ *r* ≤ −0.27,*p* < 0.001) suggest that more productive strawberry plants that produce commercial fruits with greater weight present greater speed of leaf emission (lower PHYL value) and high water use efficiency (lower values of WUE).

Five factors that presented eigenvalues greater than 1 were retained and accounted for 90.48% of the total variation among the traits (Table 2, Supplementary Table S4). Thus, it was possible to reduce the data dimensionality by ~ 78% keeping a high explanatory power. The 22 traits were grouped into the factors (FA) as follows: In FA1 the production-related traits FY, NCF, PHYL, TNF, TWF, WCF, and WUE with positive loadings, and NNCF and WNCF with negative loadings; In FA2, the traits FIRM, NDBF, NDBH, NDFF, and TSS (with negative loadings); In FA3, the pulp coloration-related traits CHROMA (with positive loading), H, and L (with negative loadings); In FA4, the traits AWNCF, TA, and TSS/TA (with negative loadings); and in FA5, the traits AWCF and OAWF. Loadings resulting from an orthogonal rotation ranges from −1 to +1 and are the correlation coefficients between each item and the factor.

**Table 2.**
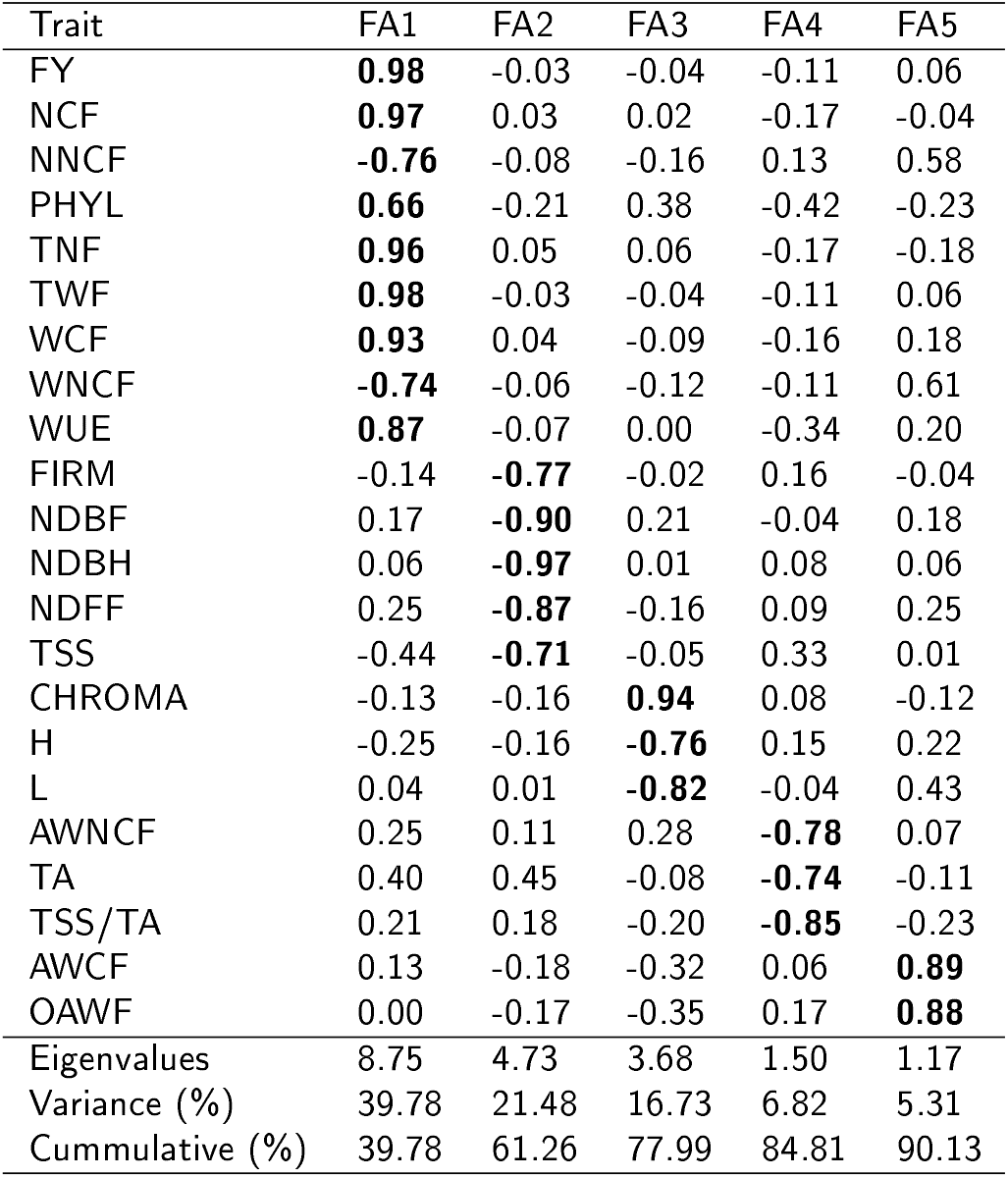
Eigenvalues, explained variance, factorial loadings after varimax rotation, and communalities obtained in the factor analysis.

**Table 3.** Selection differentials for productive, qualitative, and physiological strawberry traits. Bold values indicate traits with un-desired selection differentials.

| Trait | Factor | Goal | Xo | Xs | SD (%) |
| --- | --- | --- | --- | --- | --- |
| NNCF | FA1 | Low | 7.20 | 6.44 | -10.54 |
| WNCF | FA1 | Low | 65.36 | 62.67 | -4.11 |
| PHYL | FA1 | Low | 164.45 | 141.08 | -14.21 |
| NCF | FA1 | High | 23.47 | 24.53 | 4.53 |
| TNF | FA1 | High | 30.66 | 30.89 | 0.73 |
| WCF | FA1 | High | 358.57 | 399.60 | 11.44 |
| TWF | FA1 | High | 412.76 | 461.29 | 11.76 |
| FY | FA1 | High | 30957.11 | 34596.88 | 11.76 |
| WUE | FA1 | Low | 115.84 | 92.56 | -20.09 |
| NDBF | FA2 | Low | 57.70 | 42.25 | -26.78 |
| PF | FA2 | Low | 76.27 | 61.25 | -19.69 |
| NDBH | FA2 | Low | 77.80 | 62.38 | -19.82 |
| TSS | FA2 | High | 7.43 | 8.14 | 9.57 |
| FIRM | FA2 | High | 1.67 | 1.87 | 12.03 |
| L | FA3 | High | 54.81 | 56.11 | 2.38 |
| **CHROMA** | **FA3** | **High** | **44.60** | **44.28** | **-0.72** |
| H | FA3 | High | 43.07 | 43.12 | 0.11 |
| AWNCF | FA4 | Low | 8.73 | 8.48 | -2.87 |
| **TA** | **FA4** | **Low** | **1.42** | **1.48** | **4.39** |
| TSS/TA | FA4 | High | 5.39 | 5.57 | 3.31 |
| OAWF | FA5 | High | 13.54 | 14.97 | 10.57 |
| AWCF | FA5 | High | 14.85 | 16.31 | 9.86 |

### 3.3. Treatment ranking according to the multi-trait index

Figure 2 shows the treatment ranking according to the MGIDI index. Based on the selection pressure (~15%), the top 2 treatments selected were T12 and T16. These two treatments have a common cultivar (Albion), transplant origin (Imported), and burnt rice husk as the main component of the substrate used, differing only regarding the second component of the substrate (Organic substrate for T12 and Carolina^®^ substrate for T16 (Table 1).

**Figure 2:**
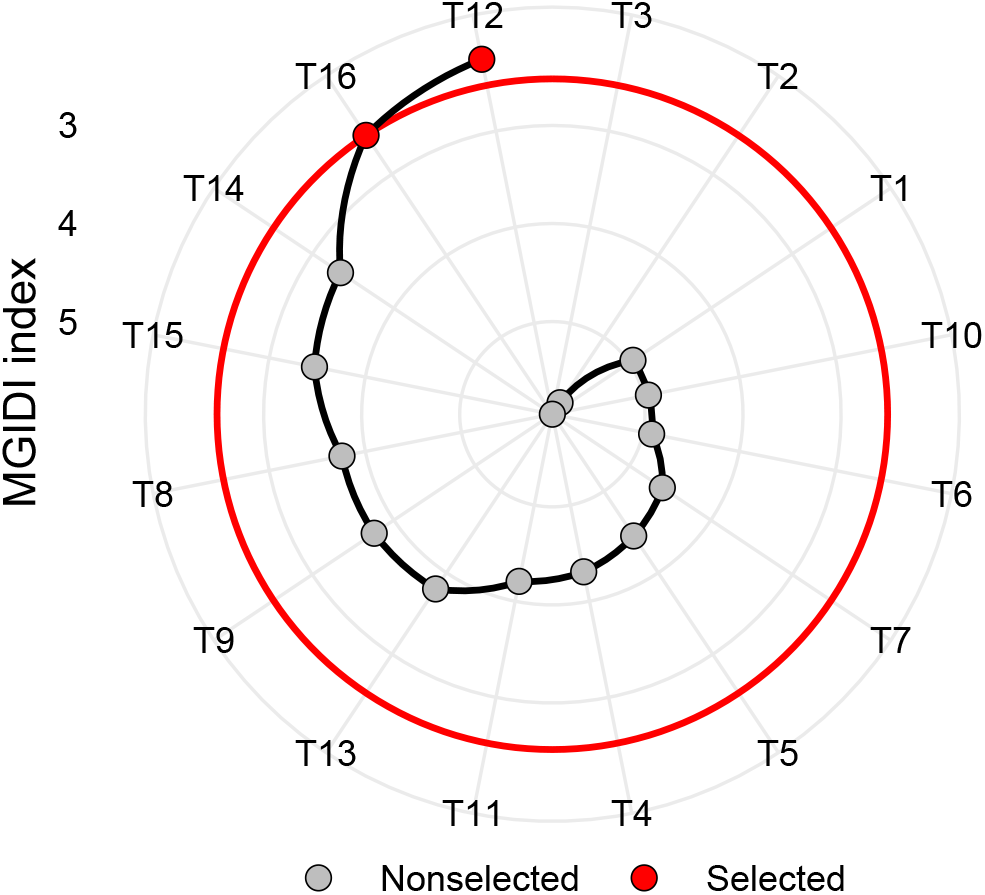
Treatment ranking based on the MGIDI index. The top 2 treatment were used to compute the selection differentials in Table 3

The MGIDI index provided desired selection differential (SD) for 20 of the 22 studied traits, which represents a success rate of ~ 91% in selecting traits with desired values. The two traits with undesired SD were Briefly, T12 and T16 provide desired values for most strawberry traits but provide fruits with higher values of titratable acidity and slightly lower chroma value, indicating a shift from a vivid to a more dull color.

The SD for traits in which high values are desired ranged from −0.72% (CHROMA) to 12.03% (FIRM) with mean SD of 6.71%. For traits in which low values are desired, the SD ranged from −26.78% (NDBF) to 4.39% (TA) with a mean SD of −12.64%.

The selection based on fruit yield solely would rank T9 and T10 as the best-performing treatments (Supplementary Fig. S16). These treatments differ from T12 and T16 mainly regarding the cultivar used (Camarosa). The direct and univariate selection, although providing higher SD for fruit yield and weight of fruits (Supplementary Table S5), is not efficient since presented desired gains for only 13 of the 22 studied traits. For example, compared to T12 and T16, T9 and T10 present on average a period 29% higher for the beginning of the harvest, 84% more non-commercial fruits, 14,5% less soluble solids, and 17.6% less flesh firmness (Tabic 3, Supplementary Table S5).

### 3.4. The strengths and weaknesses view

Figure 3 shows the strengths and weaknesses view of the treatments. The contribution of each factor to the MGIDI index is ranked from the most contributing factor (close to plot center) to the less contributing factor (close to the plot edge). The selected treatments have strengths related to FA2, which indicates that T12 and T16 present high productive precocity, i.e., lower values for NDBF, PF, NDBH (Supplementary Figs. S5-S7), higher values for TSS (Supplementary Fig. S18), and high values for FIRM (Supplementary Fig. S20).

**Figure 3.**
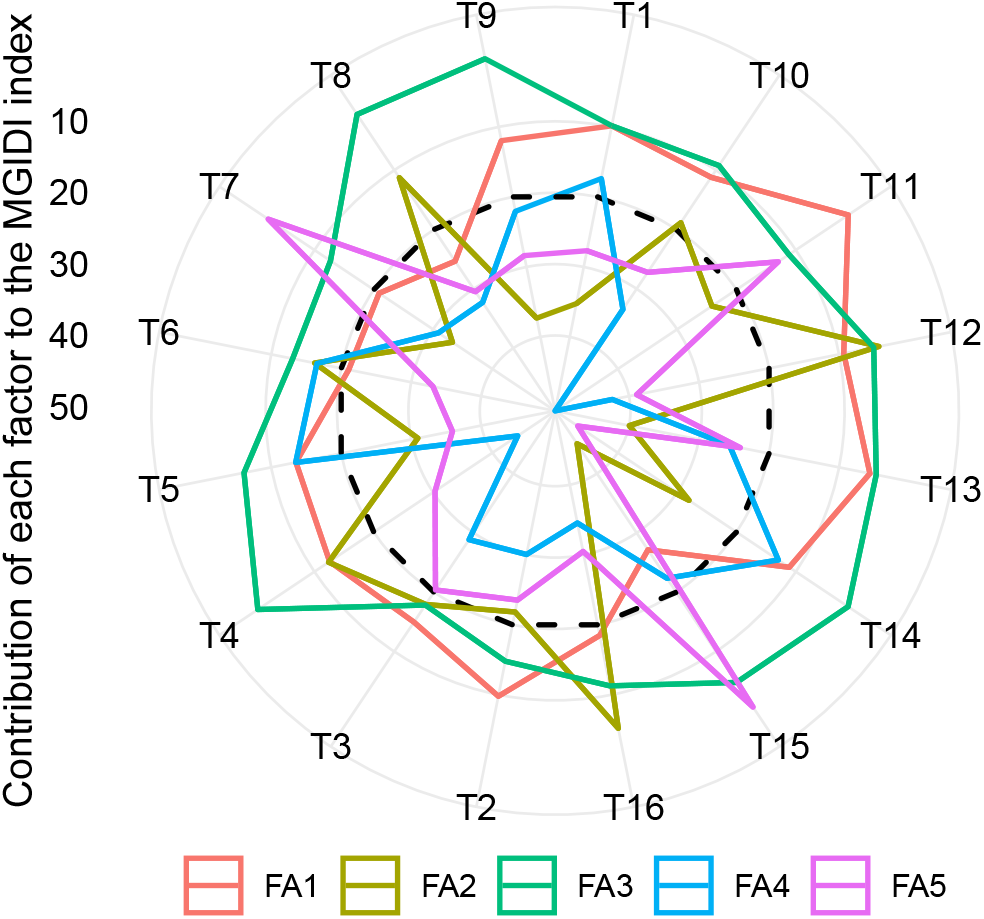
The strengths and weaknesses view of treatments is shown as the proportion of each factor on the computed multitrait genotype-ideotype distance index (MGIDI). The smallest the proportion explained by a factor (closer to the external edge), the closer the traits within that factor are to the ideotype. The dashed line shows the theoretical value if all the factors had contributed equally.

Comparing T12 and T16 regarding FA1, we conclude that T12 performed well for most of the traits within FA1, which is inferred by the lower contribution of FA1 for T12 (Fig. 3). This treatment had low phyllochron values (Supplementary Fig S8), higher values for TNF (Supplementary Fig S11), WCF (Supplementary Fig S12) TWF (Supplementary Fig S13), and FY (Supplementary Fig S16).

Both T12 and T16 have weaknesses related to FA4 (Fig. 3), which indicates that the strawberry fruits coming from these treatments presents poor values for AWNCF, TA, and TSS/TA. For example, both treatments have above-average values for TA (Supplementary Fig. S9), which is undesired in our case. It should be pointed out that it is possible to easily compare treatments using Fig. 3. For example, compared to T12 and T16, T11 presents poor performance for FA4, showing high values of AWNCF (Supplementary Fig. S4), TA (Supplementary Fig. S9), and low values of TSS/TA (Supplementary Fig. S19).

## 4. Discussion

### 4.1. Multitrait approach is more efficient than univariate for treatment recommendation

Our experimental results suggest that most of the productive, phenological, and quality traits in strawberry are influenced by the cultivar used, transplant origin, cultivation substrates, as well as by the interaction between cultivar and origin of transplants. Previous studies also reported a significant effect of cultivars for physicochemical traits of fruits (Gabriel et al., 2019), phyllochron and phenology (Diel et al., 2017) and external morphanatomy (Costa et al., 2019). This is explained because the growth and development of strawberry are regulated by the interaction of complex factors, such as temperature (Ledesma and Kawabata, 2016), light-temperature interactions (Fu et al., 2017), daylength conditions (Sønsteby et al., 2016), and cultivation substrate (Martínez et al., 2017).

As new studies evaluate a growing amount of traits aiming at better explaining the phenomena (e.g., Martínez et al., 2017; Habibzadeh et al., 2019; Sadowska et al., 2020), the challenge of summarizing the complex results into a succinct and easy-to-interpret recommendation arisen. Differently from an approach in which a *post-hoc* test, e.g., Tukey’s honest significance test is used, we have shown here how treatments in experiments with strawberry can be ranked from most desired to most undesired based on the values of several traits (Fig. 2). This approach indicated that the Albion cultivar originated from imported transplants grown in substrates where the main component (70%) is burnt rice husk provides desired values for 20 of a total of 22 productive, qualitative, physiological, and phenological traits assessed in this experiment.

Considering a univariate recommendation based on fruit yield solely, Camarosa cultivar would be recommended instead Albion since, on average, it is more productive (Supplementary Fig S16A). Previous studies also reported the high total fresh fruit mass of Camarosa compared to Albion (Castoldi da Costa et al., 2020). This is explained mainly because Camarosa can increase the emission of crowns compared to Albion, resulting in a higher number of leaves. For this reason, at the end of the cycle, the cultivar Camarosa would probably have a higher total number of leaves per plant compared to Albion (Costa et al., 2019; Diel et al., 2017) and consequently, a higher evapotranspiration rate (Martínez-Ferri et al., 2016).

The recommendation of Camarosa instead of Albion would provide undesired values for productive traits such as higher number and weight, of non-commercial fruits, phenological traits such as higher period to the beginning of the harvest, and qualitative traits such as lower values of total soluble solids and flesh firmness. Previous studies have also shown that the beginning of flowering is similar for Camarosa and Albion, but full flowering is achieved early in Albion (Castoldi da Costa et al., 2020). This results in the late beginning of harvest for Camarosa compared to Albion (Supplementary Fig. S7A).

The higher number and weight, of non-commercial fruits observed in Camarosa (Supplementary Fig. S2-S3) may result in difficulties in fruit, marketing since small fruits are not desired by consumers (Wang et al., 2017). Besides, lower irrigation water use efficiency for marketable fruit yield, i.e., a smaller mass of commercial fruits produced per liter of transpired water, can be observed (Martínez-Ferri et al., 2016).

Other important variables that can have undesired values when recommending Camarosa cultivar are flesh firmness and total soluble solids. Flesh firmness is very important since it allows identifying cultivars that produce fruits with longer shelf life and more resistant to transport damages and quality loss (Bieniasz et al., 2012). The content of total soluble solids characterizes the sweetness in fruits (Silva et al., 2015), which combined with components such as strawberry furanone defines the fruity flavors that characterize a fresh strawberry (Forney et al., 2000).

### 4.2. Strengths and weaknesses of treatments

Using the *strengths and weaknesses* view (Fig. 3) of the MGIDI index it was possible to identify the strengths and weaknesses of treatments based on a multitrait framework. The selected treatments (T12 and T16) have as strengths high productive precocity (Supplementary Figs. S5-S7), higher values for total soluble solids (Supplementary Fig. S18), and high values for flesh firmness (Supplementary Fig. S20). These strengths are mainly related to the Albion cultivar and the Imported origin of transplants. Our results corroborate with Gabriel et al. (2019), which studying 13 strawberry cultivars observed that Albion presents high values for fresh firmness and °Brix.

Previous studies have shown that Camarosa cultivar produced more acidic fruits (0.92% citric acid, lower values of TSS/TA ratio (7.7), and lower water use efficiency compared to Ventana cultivar (Neocleous, 2012). In our study, however, we observed (Supplementary Fig. S9, Supplementary Table S3) a significantly lower value for total acidy for Camarosa (1.29% citric acid). Thus, the weaknesses of T12 and T16 related to FA4 (3) are mainly related to the Albion cultivar and Imported transplant origin. It should be pointed out that fruit acidity is highly influenced by cultivars and the differentiated phenotypic behavior in relation to growing locations (Gabriel et al., 2019) or even by the different growing seasons within the same environment (Neocleous, 2012).

Even though T12 and T16 had provided desired selection differentials for all traits within FA1, special attention should be given to the traits within this factor, and improving these characteristics is a crucial point in looking for an *ideal* treatment. For example, the water use efficiency was highly influenced by the cultivation substrate and less by the cultivar used (Supplementary Table S3; Supplementary Fig. S17). The high water use efficiency in substrates with burnt rice husks (Supplementary Fig. S1C) seems to be related to the higher water retention capacity at lower tensions (Fig. 1), which can be explained by its favorable physical proprieties such as lower total pososity and aeration space, resulting in higher amounts of readily available water (Supplementary Table S2). These physical characteristics are crucial for root growth and plant development due to its promotion of an adequate air/water balance during plant cultivation (Terra et al., 2011).

Our results provide pieces of evidence that Albion was less efficient in water use than Camarosa (higher values of WUE). However, this diverges from Neocleous (2012) that observed a larger amount of water consumed to produce one kg of fruit fresh weight for the Camarosa cultivar and from Martínez-Ferri et al. (2016), which have shown that Camarosa is the less-efficient cultivar producing 7.6 g of marketable fruits per liter of water used, much less than Fortuna cultivar, which produced 26 g of marketable fruits per liter of water used.

Strawberry requires a high amount of water and although cultivars with more leaves can produce a higher amount of fruits, a higher total leaf area per plant can lead to higher transpiration rates, increasing the water consumption per fruit produced. It should be pointed out that the water use efficiency in this study was computed considering the total fruit production; so the water use efficiency of Camarosa considering the production of marketable fruits would possibly be smaller due to its higher weight of noncommercial fruits produced compared to Albion (Supplementary Fig. S3).

Since Albion presents interesting quality features (Fig. 3), Breeders and Agronomist efforts should be focused on increasing fruit production of Albion cultivar as well as its water use efficiency in burnt rice husk-based substrates. This would be achieved by (i) increasing the uptake efficiency of available water through the plant-system, e.g., by improving root volume and surface area Costa et al. (2019); (ii) improving crop transpiration efficiency by acquiring more carbon per water transpired, e.g., by improving the control of stomatal closure aiming at a positive impact on daily vapor pressure deficit (Sinclair, 2018); and (iii) increasing the partitioning of the acquired biomass into the harvested product, e.g., by a better understanding of patterns of plant biomass partitioning depending on nitrogen source (Cambui et al., 2011).

### 4.3. Where are we now and where can we go?

Horticultural experiments that assess multiple traits are common but the benefits of a multi-trait framework analysis seem to be not fully exploited. For example, the study by Martínez et al. (2017) evaluated 12 yicld- and quality-related traits in strawberries grown in different growth mediums. Another example is the study by Aaby et al. (2012) which evaluated phenolic compounds in fruits of 27 strawberry cultivars. For both examples, the inferences were made based on a univariate approach using a post hoc test for multiple mean comparisons. While this is methodologically correct, the inferences can be limited because the correlation structure of the data is not taken into account. In addition, the ambiguity in multiple comparisons can confound more than clarify the differences among treatments/cultivars. For example, when comparing the cultivars for the phenolic compounds using the Tukey’s test, Aaby et al. (2012) have observed means followed by “cdefghij”. This may not be the simplest way to identify the best cultivar for one trait, and for sure is not the better strategy for recommending a cultivar based on information of multiple traits.

While multivariate approaches have been used with the main goal of investigating the interrelationships between traits and grouping of the variables in a PCA plot (e.g., González-Domínguez et al., 2020; Šamec et al., 2016), its use in the analysis of strawberry experiments seems to be more an exception than a rule. A new approach that applies principal component analysis on coefficients of nonlinear growth models was introduced by Diel et al. (2020). In this approach, the use of growth models substantially increases the inferences that can be made about crop growth (fruit production), and the PCA summarizes these pieces of information, simplifying its interpretation. This approach presents an important advance over existing literature but can be applied in one trait at once.

The frontier of the MGIDI index in the evaluation of horticultural crops is expected to rapidly expand. The key point underlying its use is choosing an *ideal* treatment, which will certainly vary depending on the goal of the study. In our example, we were able to rank the treatments from the most desired to the most undesired, based on information on multiple traits (Fig. 2), and tthe strengths and weaknesses view of treatments (Fig. 3) stand out as a powerful tool to develop better recommendation strategies. Using the MGIDI index in future studies will dramatically reduce the number of tables/figures needed, making it easier for the recommendation of superior treatments in experiments with strawberries

## 5. Conclusions

Using the multi-trait genotype-ideotype distance index (MGIDI), we have shown how to select superior treatments in experiments with strawberry where multiple traits have been assessed. In our study, the treatments selected by the MGIDI index were characterized by the Albion cultivar originated from imported transplants grown in substrates where the main component (70%) is burnt rice husk. The selected treatments provided desired values for 20 from a total of 23 productive, qualitative, physiological, and phenological evaluated traits. The *strengths and weakness view* provided by the MGIDI index revealed that looking for an ideal treatment should direct the efforts on improving the water efficiency use and reducing total acidy of fruits. On the other hand, the strengths of selected treatments are mainly related to productive precocity, total soluble solids, and flesh firmness. Overall, this study provides new insights into how the MGIDI index can provide a multivariate framework for the analysis of experiments with strawberries. The MGIDI index stands out as a powerful tool to develop better recommendation strategies, thus contributing to the environmental social, and economic framework of strawberry cultivation systems.

### CRediT authorship contribution statement

**Tiago Olivoto:** Conceptualization, Methodology, Software, Formal analysis, Writing - Original Draft. **Maria Inês Diel:** Data curation, Supervision, Investigation, Writing - Original Draft. **Denise Schmidt:** Supervision, Investigation, Writing - Review & Editing. **Alessandro Dal’Col Lúcio:** Supervision, Investigation, Writing - Review & Editing.

## Declaration of Competing Interest

The authors declare that they have no known competing financial interests or personal relationships that could have appeared to influence the work reported in this paper.

## Acknowledgements

We thank the National Council for Scientific and Technological Development (CNPq) and Coordination for the Improvement of Higher Education Personnel (CAPES) for fellowships and grants to the authors

## Notes

### Competing Interest Statement

The authors have declared no competing interest.

