## Supplementary for "Multivariate analysis of strawberry experiments: where are we now and where can we go?"

### Supporting Informatin for Uncovering the secrets behind the multivariate analysis of strawberry experiments by

**3 Appendix C - Supplementary tables** **32**

### Getting started

This supplementary material has three main sections. In the first one (Appendix A), we present the R datasets, the packages, and all R codes needed to reproduce the examples described in the paper. The second section (Appendix B), contains the supplementary figures, and the last one (Appendix C) contains the supplementary tables.

### 1 Appendix A - R codes

#### 1.1 R packages

To reproduce the examples of this material, the R packages `metan`, `ggplot2`, `rio`, and `patchwork` are needed.

```
library(metan) # Mixed model and mtsi index
library(ggplot2)
library(dplyr)
library(patchwork)
```

For reasons of confidentiality, only the structure of the data (`df`) is shown. One can, of course, adapting the codes we provided here to an own data set, provided that the structure of the data is kept.

```
str(df)
# tibble [64 x 27] (S3: tbl_df/tbl/data.frame)
#  $ REP   : Factor w/ 4 levels "1","2","3","4": 1 1 1 1 1 1 1 1 1 1 ...
#  $ SUB   : Factor w/ 4 levels "BC_CO","BC_SC",..: 1 1 1 1 2 2 2 2 3 3 ...
#  $ CUL   : Factor w/ 2 levels "ALB","CAM": 2 2 1 1 2 2 1 1 2 2 ...
#  $ ORI   : Factor w/ 2 levels "IMP","NAC": 2 1 2 1 2 1 2 1 2 1 ...
#  $ TRAT  : Factor w/ 16 levels "T1","T10","T11",..: 1 9 10 11 12 13 14 15 16 2 ...
#  $ NNCF  : num [1:64] 3.88 6.12 1.2 2.86 4.67 ...
#  $ WNCF  : num [1:64] 21.4 36.2 10.3 50.1 41.1 ...
#  $ AWNCF : num [1:64] 5.51 5.92 8.62 17.55 8.81 ...
#  $ WUE   : num [1:64] 93.4 125.5 136.3 94.2 190.5 ...
#  $ NDBF  : num [1:64] 82 61 69 44 86 61 71 44 54 55 ...
#  $ NDFF  : num [1:64] 92 83 82 70 92 89 82 70 69 70 ...
#  $ NDBH  : num [1:64] 81 82 78 69 108 85 88 72 74 79 ...
#  $ PHYL  : num [1:64] 162 185 189 180 124 ...
#  $ TA    : num [1:64] 1.38 1.72 1.77 1.4 1.51 ...
#  $ NCF   : num [1:64] 28.1 24 19.6 24.9 12.3 ...
#  $ TNF   : num [1:64] 32 30.1 20.8 27.7 17 ...
#  $ WCF   : num [1:64] 425 307 330 372 148 ...
#  $ TWF   : num [1:64] 446 343 340 423 189 ...
#  $ AWCF  : num [1:64] 15.1 12.8 16.8 15 12 ...
#  $ OAWF  : num [1:64] 13.9 11.4 16.4 15.3 11.1 ...
#  $ FY    : num [1:64] 33473 25713 25509 31698 14170 ...
#  $ TSS   : num [1:64] 6.77 8.27 7.25 8.35 7.2 ...
#  $ TSS/TA: num [1:64] 4.89 4.79 4.1 5.98 4.76 ...
#  $ FIRM  : num [1:64] 1.53 1.95 2.07 1.93 1.73 ...
#  $ L     : num [1:64] 57 48.7 55.2 53.2 53.4 ...
#  $ CHROMA: num [1:64] 40.5 47.1 46 47 43.7 ...
#  $ H     : num [1:64] 42.4 40.7 43.2 42.7 42.2 ...
```

#### 1.2 Analysis of Variance

The function `compute_anova()` computes a three-way ANOVA for each variable in `df`. The function `get_anova_table()` get the ANOVA table and `get_predicted_val()`get the predicted values.

```
compute_anova <- function(df,
                          rep,
                          sub,
                          cul,
```

```
                         ori,
                         resp){
  factors <- df %>% select_cols({{rep}}, {{sub}}, {{cul}}, {{ori}})
  vars <- df %>% select({{resp}}, -names(factors)) %>% select_numeric_cols()
  nvar <- ncol(vars)
  model_formula <- paste("Y ~ REP + SUB*ORI*CUL")
  listres <- list()
  vin <- 0
  for (var in 1:nvar) {
    data <- factors %>% mutate(Y = vars[[var]])
    listres[[paste(names(vars[var]))]] <- lm(model_formula, data = data)
  }
  return(listres)
}
get_anova_table <- function(models){
  models %>% map_dfr(~.x %>% anova() %>% tidy(), .id = 'TRAIT')
}
get_predicted_val <- function(models){
  predicted <- models %>% map_dfc(~.x %>% predict())
  levels <- models[[1]][["model"]] %>% select_non_numeric_cols()
  df <- cbind(levels, predicted)
  return(df)
}

# Fit the models
models <- compute_anova(df, REP, SUB, CUL, ORI, everything())

# Get anova table
anovas <- get_anova_table(models)
```

#### 1.3 The MGIDI index

The MGIDI index is computed with the function `mgidi()`. First, we need to create a two-way table with the predicted values for each treatment in rows and traits in columns.

```
pred_vals <- get_predicted_val(models)
df_mgidi <-
  pred_vals %>%
  remove_cols(REP) %>%
  add_cols(TRAT = rep(paste("T", 1:16, sep = ""), length.out = 64), .after = CUL) %>%
  mutate(CUL = fct_relevel(CUL, rev)) %>%
  mutate(ORI = fct_relevel(ORI, rev))

df_means <-
```

```
  df_mgidi %>%
  remove_cols(SUB:CUL) %>%
  means_by(TRAT) %>%
  column_to_rownames("TRAT")
round_cols(df_means, digits = 2)
#       NNCF   WNCF AWNCF    WUE  NDBF  NDFF  NDBH   PHYL   TA   NCF   TNF    WCF
# T1    7.40  69.46  9.26 100.43 79.50 94.25 89.00 184.27 1.08 23.93 31.33 333.89
# T10  12.12 106.21  8.54  62.01 55.50 74.00 76.00 136.09 1.45 39.34 51.46 542.53
# T11   5.33  56.58  9.40  86.31 57.50 71.50 75.25 175.23 1.65 28.15 33.48 468.86
# T12   6.87  65.48  8.80  78.27 42.50 61.00 62.50 129.55 1.43 27.03 33.90 442.10
# T13   9.72  84.12  8.17  72.57 61.50 80.75 85.25 153.84 1.12 31.17 40.88 643.67
# T14   8.96  73.19  7.74  85.04 56.00 76.50 73.25 122.14 1.17 27.40 36.35 379.50
# T15   2.51  22.85  8.04 113.63 62.25 77.00 83.75 197.36 1.26 19.67 22.17 318.22
# T16   6.01  59.86  8.17 106.85 42.00 61.50 62.25 152.61 1.53 22.04 27.87 357.09
# T2    7.77  81.63  9.85 161.59 64.75 86.50 82.00 146.13 1.58 18.24 26.01 237.30
# T3    4.80  56.93 10.44 204.62 69.00 76.25 82.75 224.94 1.69 12.99 17.79 202.14
# T4    7.24  66.18 10.70 117.09 46.00 62.25 64.00 200.56 1.58 19.61 26.85 293.99
# T5    8.67  70.57  8.00 118.09 69.75 86.25 87.75 164.38 1.21 22.83 31.51 314.12
# T6    7.30  60.06  7.59 180.33 56.25 84.50 80.50 165.19 1.53 15.45 22.75 184.31
# T7    3.66  38.97  8.63 153.14 62.25 86.50 86.50 196.70 1.61 14.36 18.02 240.28
# T8    5.26  47.31  9.20 152.09 44.50 68.50 68.25 172.73 1.65 16.22 21.48 236.74
# T9   11.63  86.29  7.25  61.30 54.00 73.00 85.75 109.56 1.17 37.11 48.74 542.32
#         TWF  AWCF  OAWF       FY  TSS TSS/TA FIRM     L CHROMA     H
# T1   403.34 13.84 12.91 30250.76 6.56   6.23 1.61 56.35  42.94 43.17
# T10  648.74 13.86 12.64 48655.77 7.40   5.11 1.68 51.69  46.35 41.61
# T11  525.44 16.60 15.45 39407.92 7.45   4.52 1.82 57.29  43.10 45.41
# T12  507.58 16.34 14.98 38068.55 8.25   5.85 1.87 54.31  45.54 42.89
# T13  551.16 15.97 13.60 41336.74 6.64   5.91 1.59 56.40  42.71 42.12
# T14  452.69 14.05 12.59 33951.70 7.11   6.28 1.59 51.26  45.24 44.36
# T15  341.07 16.24 15.38 25580.39 6.78   5.42 1.62 56.62  44.71 43.27
# T16  415.00 16.28 14.96 31125.20 8.02   5.29 1.87 57.91  43.02 43.35
# T2   318.93 12.73 12.08 23919.49 7.25   4.62 1.63 49.80  48.76 41.43
# T3   259.08 15.13 14.38 19430.72 8.18   4.88 1.62 58.42  39.85 45.90
# T4   360.17 14.95 13.40 27012.83 7.59   4.92 1.79 54.44  45.67 42.64
# T5   384.69 13.48 12.04 28851.77 7.21   6.26 1.61 55.21  43.43 42.74
# T6   244.37 11.90 10.65 18327.90 7.74   5.22 1.81 50.76  47.36 41.56
# T7   279.25 17.00 15.40 20943.94 7.80   4.93 1.51 53.71  46.12 42.74
# T8   284.06 14.54 13.26 21304.14 8.35   5.09 1.69 55.82  45.70 43.95
# T9   628.61 14.64 12.90 47145.92 6.53   5.70 1.40 56.92  43.07 42.05

# Compute the MGIDI index
mgidi_ind <- mgidi(df_means,
                   ideotype = c(rep("l", 9), rep("h", 13)))
#
```

```
# ---------------------------------------------------------------------------------
# Principal Component Analysis
# ---------------------------------------------------------------------------------
# # A tibble: 22 x 4
#    PC    Eigenvalues `Variance (%)` `Cum. variance (%)`
#    <chr>       <dbl>          <dbl>                <dbl>
#  1 PC1          8.75           39.8                 39.8
#  2 PC2          4.73           21.5                 61.3
#  3 PC3          3.68           16.7                 78.0
#  4 PC4          1.5             6.82                84.8
#  5 PC5          1.17            5.31                90.1
#  6 PC6          0.64            2.93                93.1
#  7 PC7          0.45            2.07                95.1
#  8 PC8          0.42            1.89                97.0
#  9 PC9          0.31            1.4                 98.4
# 10 PC10         0.13            0.59                99.0
# # ... with 12 more rows
# ---------------------------------------------------------------------------------
# Factor Analysis - factorial loadings after rotation-
# ---------------------------------------------------------------------------------
# # A tibble: 22 x 8
#    VAR     FA1   FA2   FA3   FA4    FA5 Communality Uniquenesses
#    <chr> <dbl> <dbl> <dbl> <dbl>  <dbl>       <dbl>        <dbl>
#  1 NNCF  -0.76 -0.08 -0.16  0.13  0.580        0.97         0.03
#  2 WNCF  -0.74 -0.06 -0.12 -0.11  0.61         0.95         0.05
#  3 AWNCF  0.25  0.11  0.28 -0.78  0.07         0.77         0.23
#  4 WUE    0.87 -0.07  0    -0.34  0.2          0.93         0.07
#  5 NDBF   0.17 -0.9   0.21 -0.04  0.18         0.92         0.08
#  6 NDFF   0.25 -0.87 -0.16  0.09  0.25         0.91         0.09
#  7 NDBH   0.06 -0.97  0.01  0.08  0.06         0.95         0.05
#  8 PHYL   0.66 -0.21  0.38 -0.42 -0.23         0.84         0.16
#  9 TA     0.4   0.45 -0.08 -0.74 -0.11         0.93         0.07
# 10 NCF    0.97  0.03  0.02 -0.17 -0.04         0.98         0.02
# # ... with 12 more rows
# ---------------------------------------------------------------------------------
# Comunalit Mean: 0.9012502
# ---------------------------------------------------------------------------------
# Selection differential
# ---------------------------------------------------------------------------------
# # A tibble: 22 x 8
#    VAR   Factor       Xo       Xs       SD  SDperc sense     goal
#    <chr> <chr>     <dbl>    <dbl>    <dbl>   <dbl> <chr>    <dbl>
#  1 NNCF  FA1        7.20     6.44   -0.759 -10.5   decrease   100
#  2 WNCF  FA1       65.4     62.7    -2.69   -4.11  decrease   100
```

```
#  3 WUE   FA1      116.      92.6   -23.3   -20.1   decrease   100
#  4 PHYL  FA1      164.     141.    -23.4   -14.2   decrease   100
#  5 NCF   FA1       23.5     24.5     1.06    4.53  increase   100
#  6 TNF   FA1       30.7     30.9     0.225   0.735 increase   100
#  7 WCF   FA1      359.     400.     41.0    11.4   increase   100
#  8 TWF   FA1      413.     461.     48.5    11.8   increase   100
#  9 FY    FA1    30957.   34597.   3640.     11.8   increase   100
# 10 NDBF  FA2       57.7     42.2   -15.5   -26.8   decrease   100
# # ... with 12 more rows
# --------------------------------------------------------------------------------
# Selected genotypes
# ---------------------------------------------------------------------------------
# T12 T16
# ---------------------------------------------------------------------------------
```

#### 1.4 The strengths and weaknesses view

In the following code we obtain the contribution of each factor on the MGIDI value of all treatments.

```
plot(mgidi_ind,
     type = "contribution", # Get the proportion plot
     genotypes = "all") # All hybrids (selected hybrids are plotted by default)
```

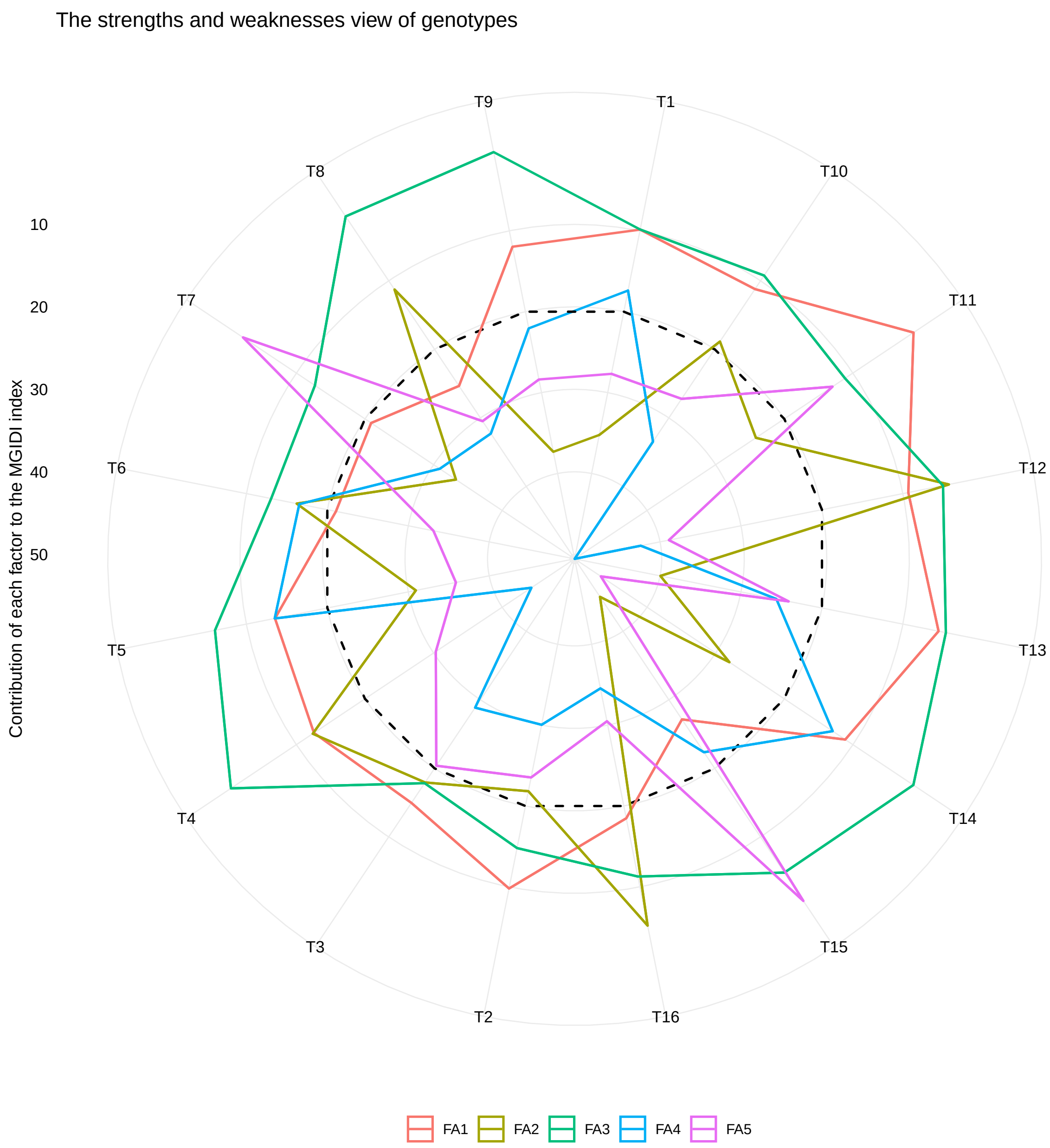

The ranks for each factor for both selected and all genotypes can be obtained by calling

```
gmd(mgidi_ind, "contri_fac_rank") # Ranks for all genotypes
# Class of the model: mgidi
# Variable extracted: contri_fac_rank
# # A tibble: 16 x 5
#    FA1   FA2   FA3   FA4   FA5
#    <chr> <chr> <chr> <chr> <chr>
#  1 T11   T12   T9    T14   T15
#  2 T13   T16   T4    T5    T7
#  3 T12   T8    T8    T6    T11
```

```
#   4 T2    T4    T14   T1    T3
#   5 T1    T6    T13   T9    T2
#   6 T14   T3    T15   T15   T13
#   7 T10   T10   T12   T13   T10
#   8 T9    T2    T5    T3    T1
#   9 T4    T11   T10   T2    T9
#  10 T5    T14   T1    T7    T4
#  11 T3    T5    T11   T8    T16
#  12 T16   T7    T16   T10   T8
#  13 T7    T1    T7    T16   T6
#  14 T6    T9    T6    T12   T5
#  15 T8    T13   T2    T4    T12
#  16 T15   T15   T3    T11   T14
```

### 2 Appendix B - Supplementary figures

#### 2.1 Correlation coefficient

```
correl <- corr_coef(df)
plot(correl, signif = "pval", size.text.signif = 2)
```

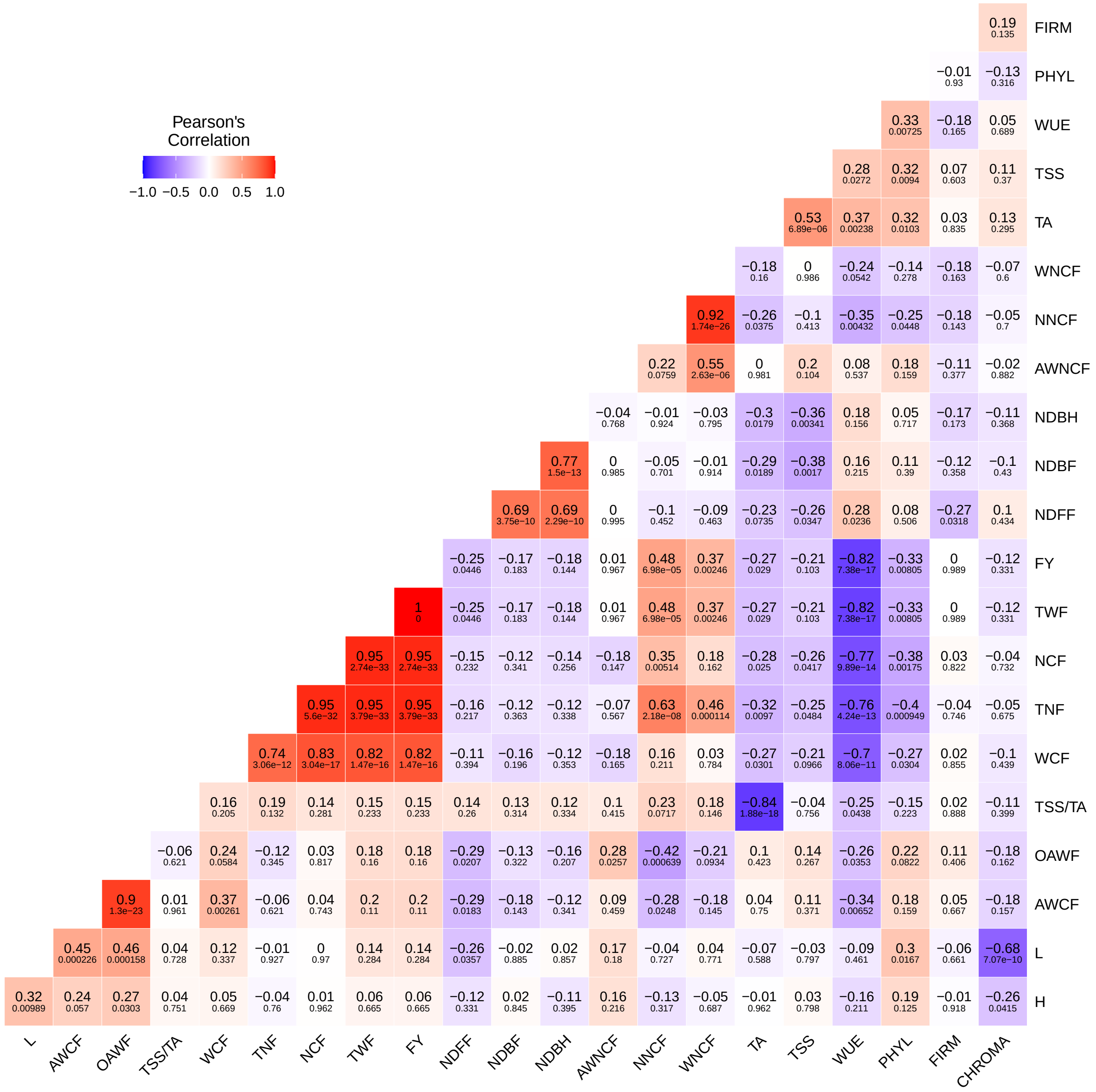

Figure S 1: Phenotypic correlation coefficient and p-values between 23 productive, qualitative and physiological strawberry traits. N = 64.

#### 2.2 Traits with negative desired gains

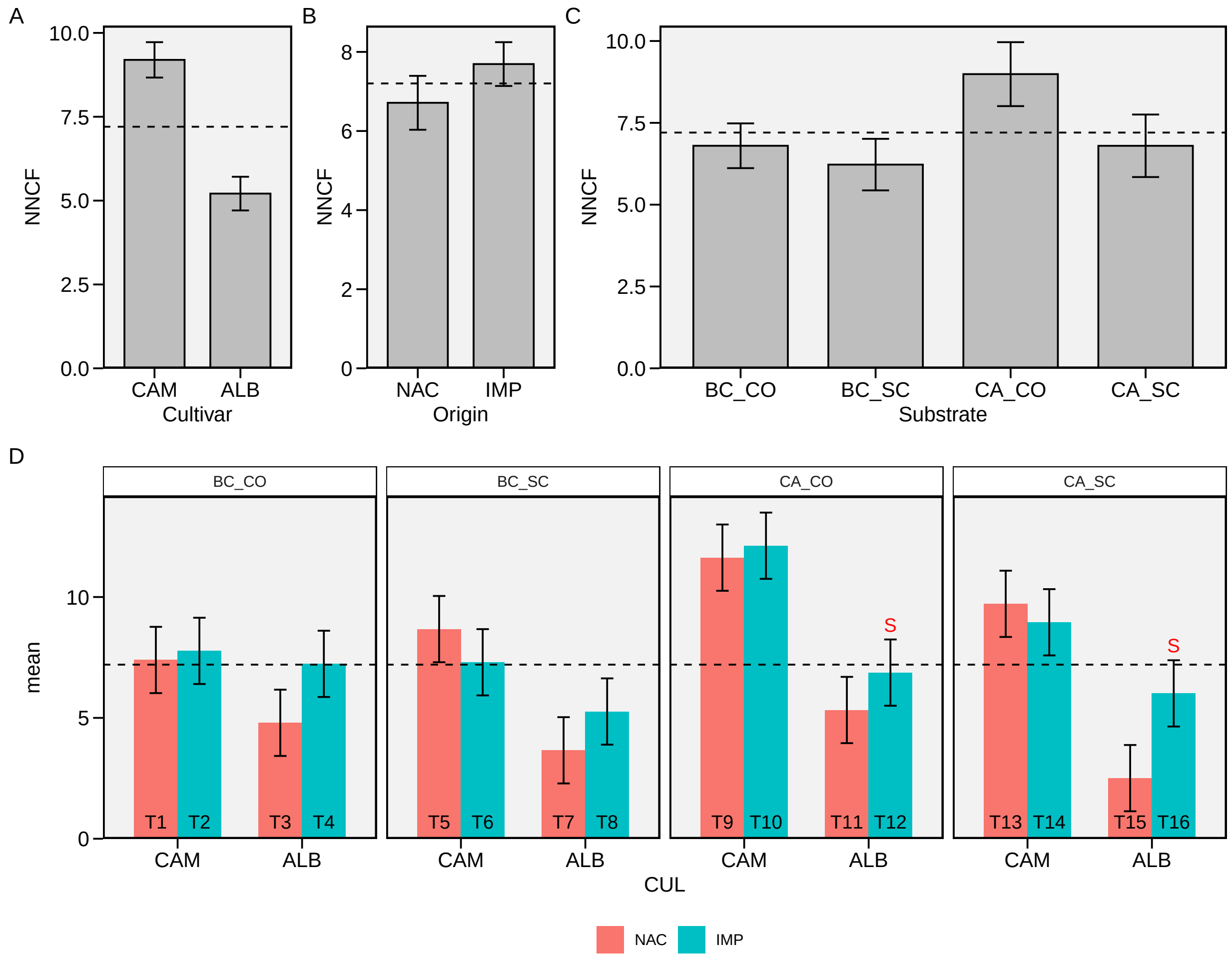

Figure S 2: Number of non-commercial fruits. Main effect of cultivar (A), origin (B), substrate (C) and for the three-way interaction (D). The horizonal dashed line shows the overall mean. The 'S' letter indicates the selected treatments. Bars shows the mean and standard error. N = 4.

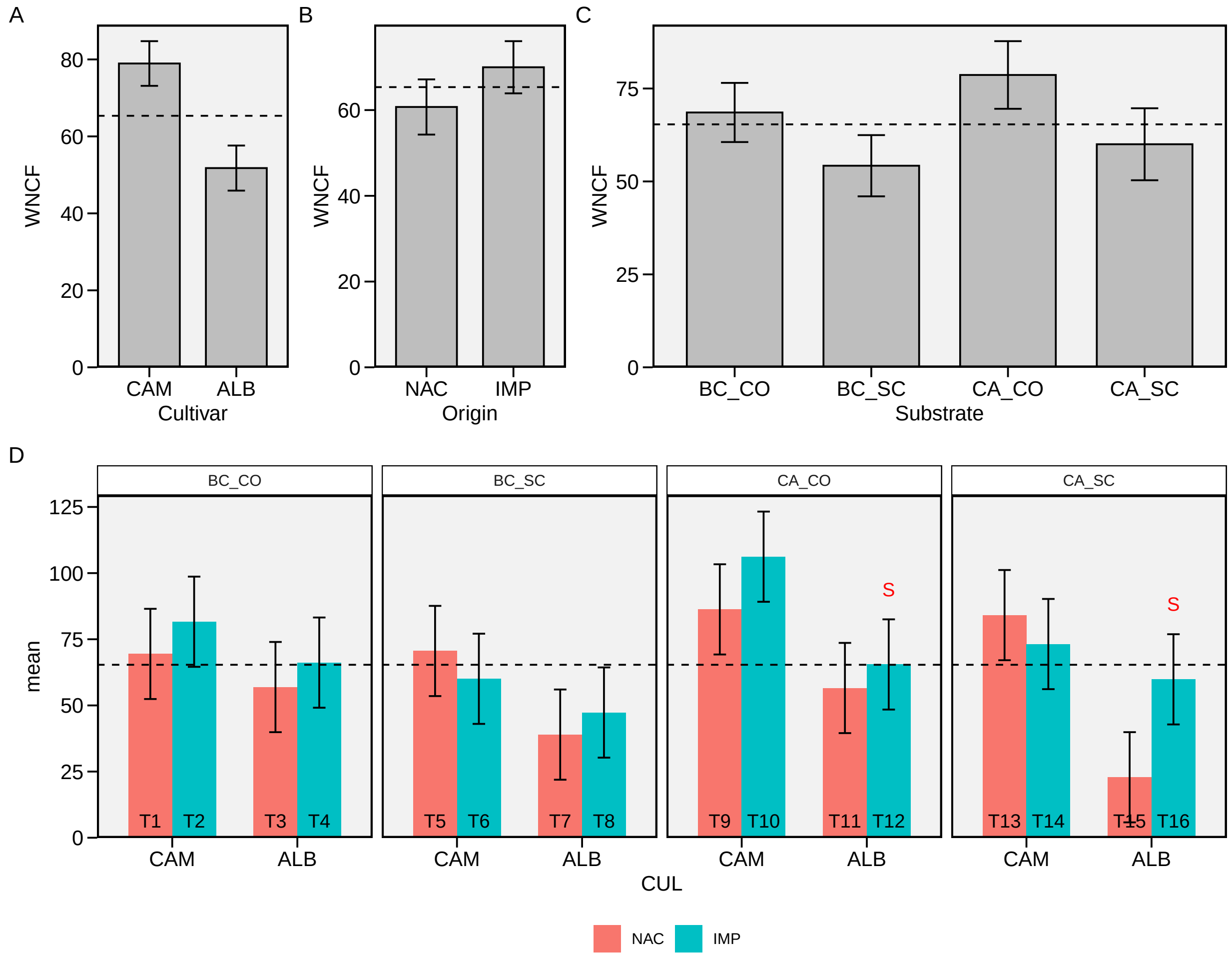

Figure S 3: Weight of non-commercial fruits. Main effect of cultivar (A), origin (B), substrate (C) and for the three-way interaction (D). The horizonal dashed line shows the overall mean. The 'S' letter indicates the selected treatments. Bars shows the mean and standard error. N = 4.

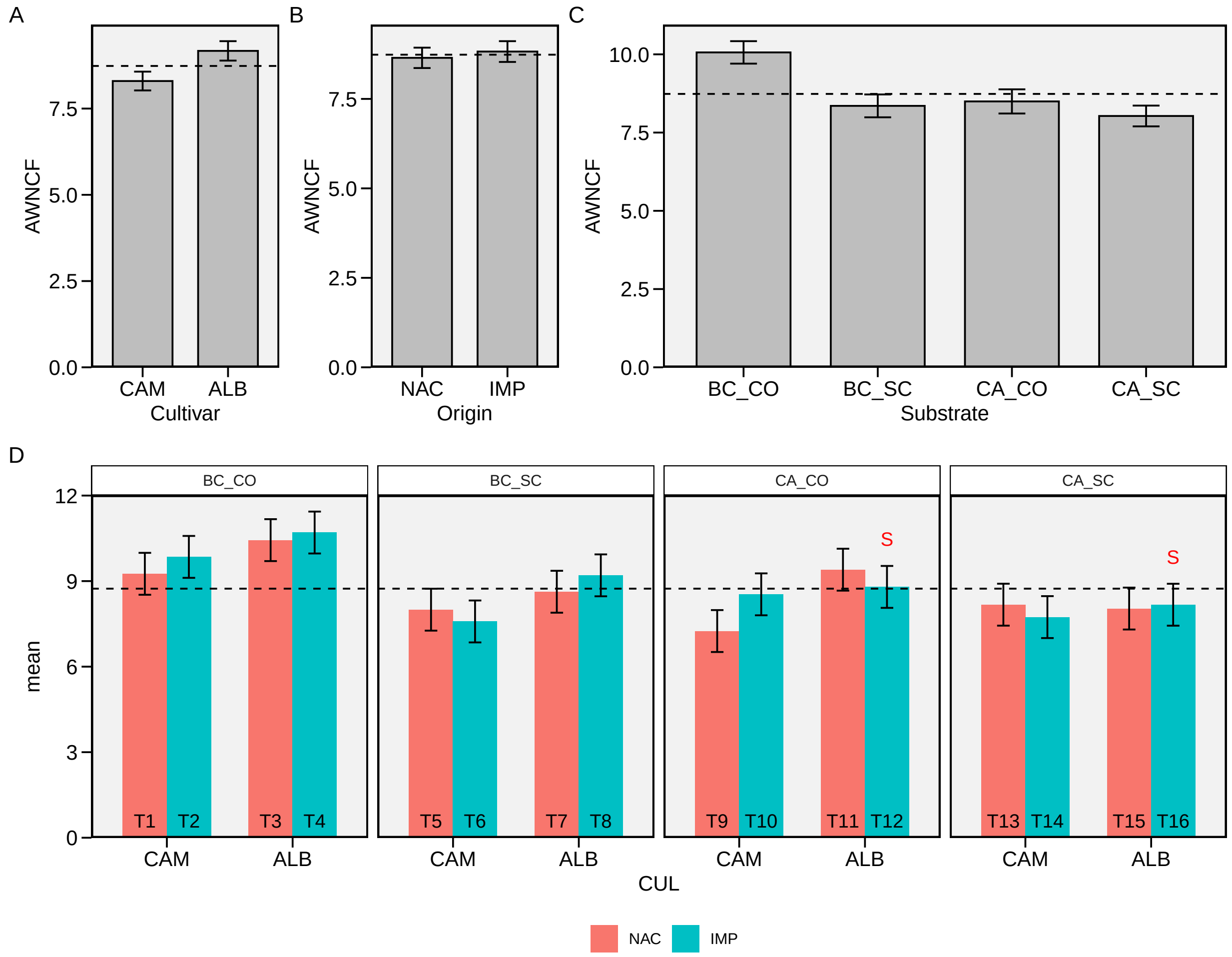

Figure S 4: Average weight of non-commercial fruits. Main effect of cultivar (A), origin (B), substrate (C) and for the three-way interaction (D). The horizonal dashed line shows the overall mean. The 'S' letter indicates the selected treatments. Bars shows the mean and standard error. N = 4.

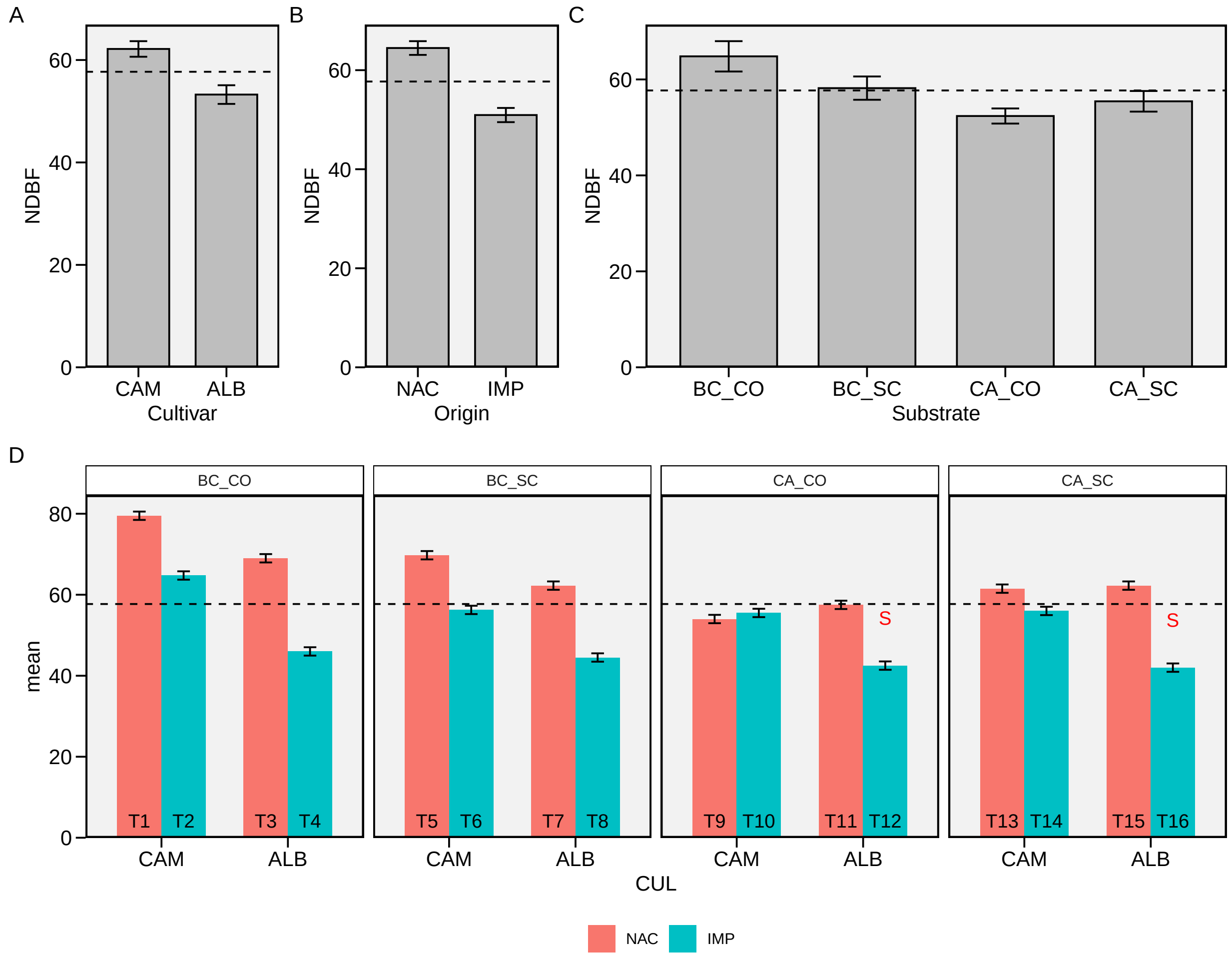

Figure S 5: Begin of floration (days). Main effect of cultivar (A), origin (B), substrate (C) and for the three-way interaction (D). The horizonal dashed line shows the overall mean. The 'S' letter indicates the selected treatments. Bars shows the mean and standard error. N = 4.

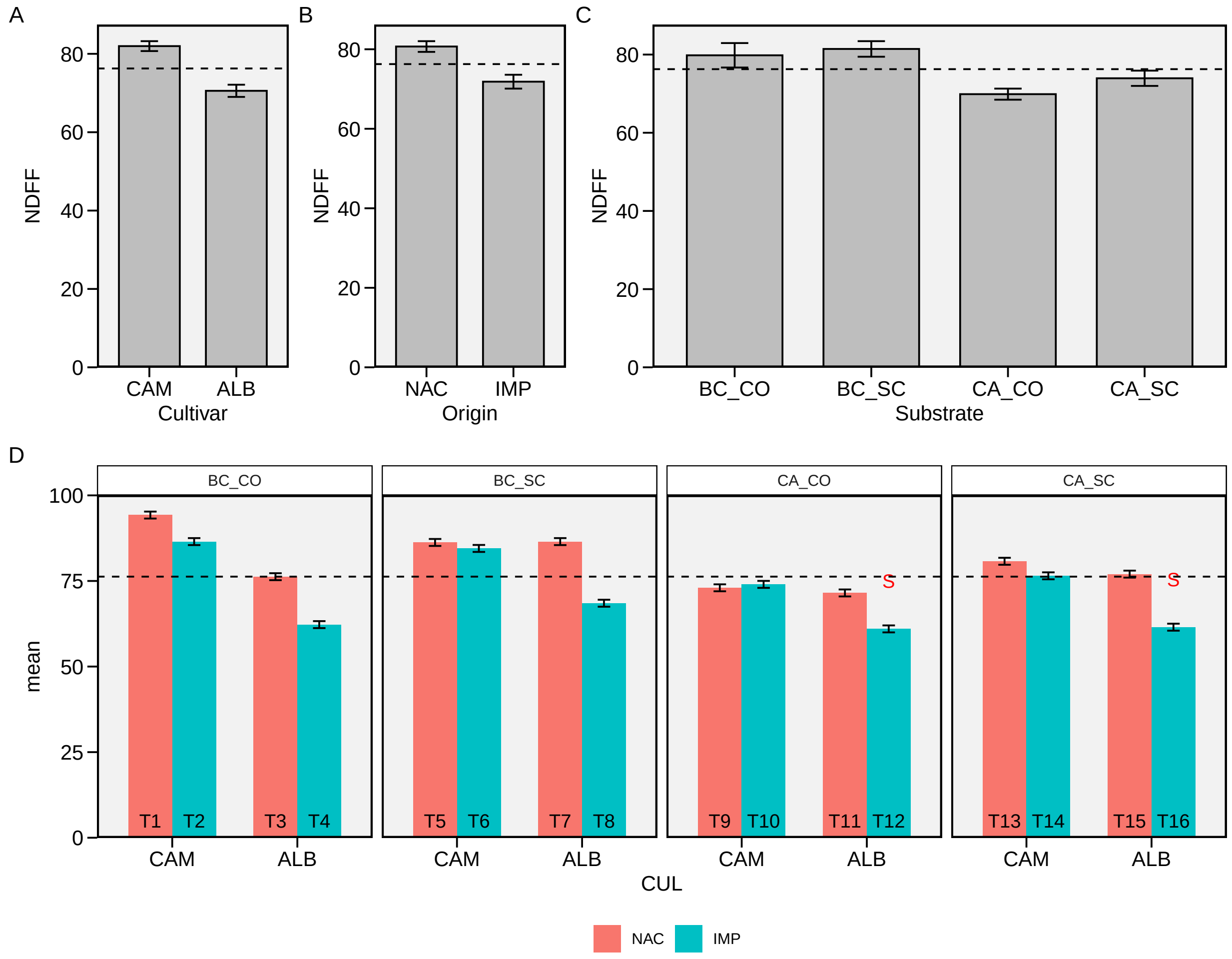

Figure S 6: Plene floration (days). Main effect of cultivar (A), origin (B), substrate (C) and for the three-way interaction (D). The horizonal dashed line shows the overall mean. The 'S' letter indicates the selected treatments. Bars shows the mean and standard error. N = 4.

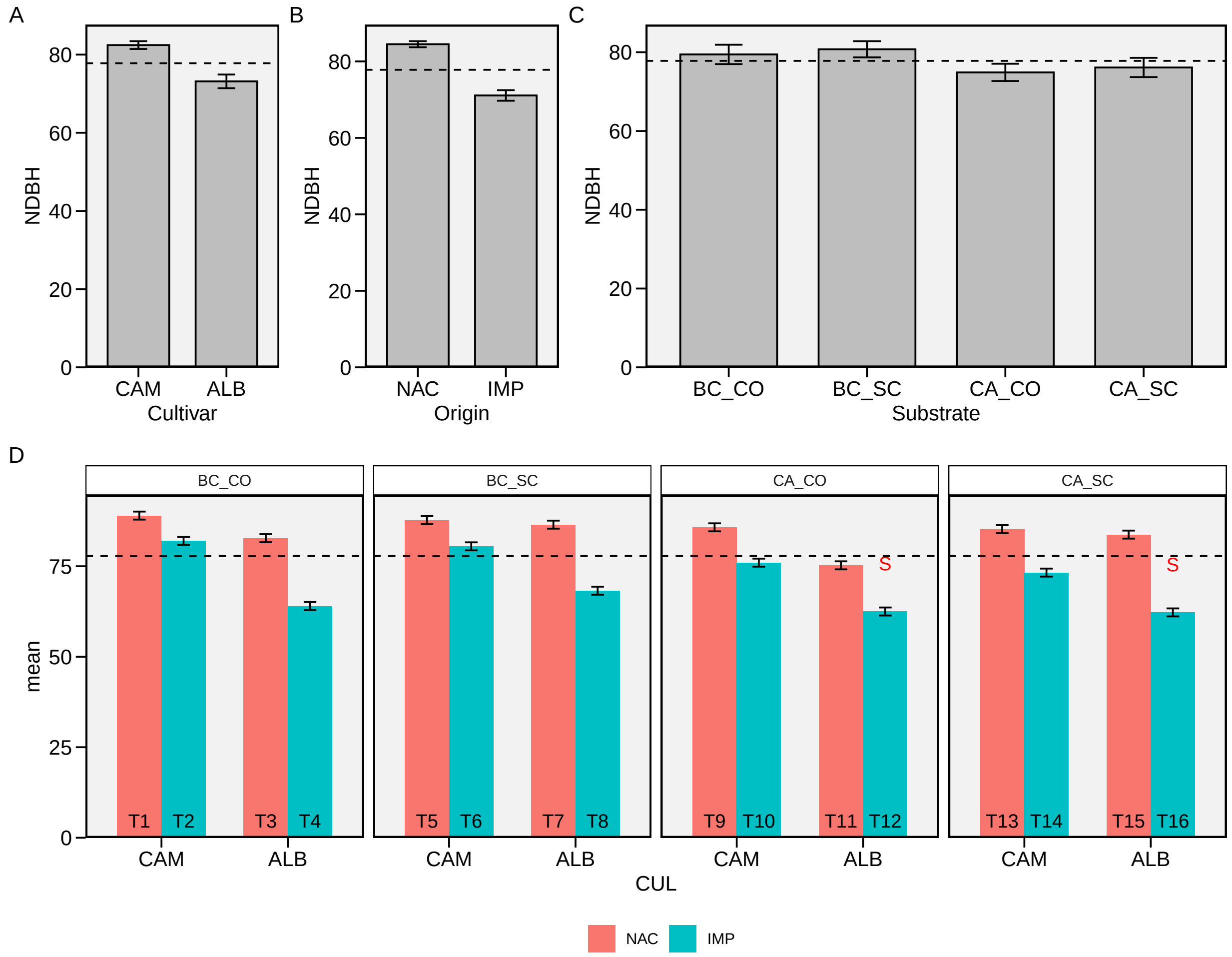

Figure S 7: Begin of harvest (days). Main effect of cultivar (A), origin (B), substrate (C) and for the three-way interaction (D). The horizonal dashed line shows the overall mean. The 'S' letter indicates the selected treatments. Bars shows the mean and standard error. N = 4.

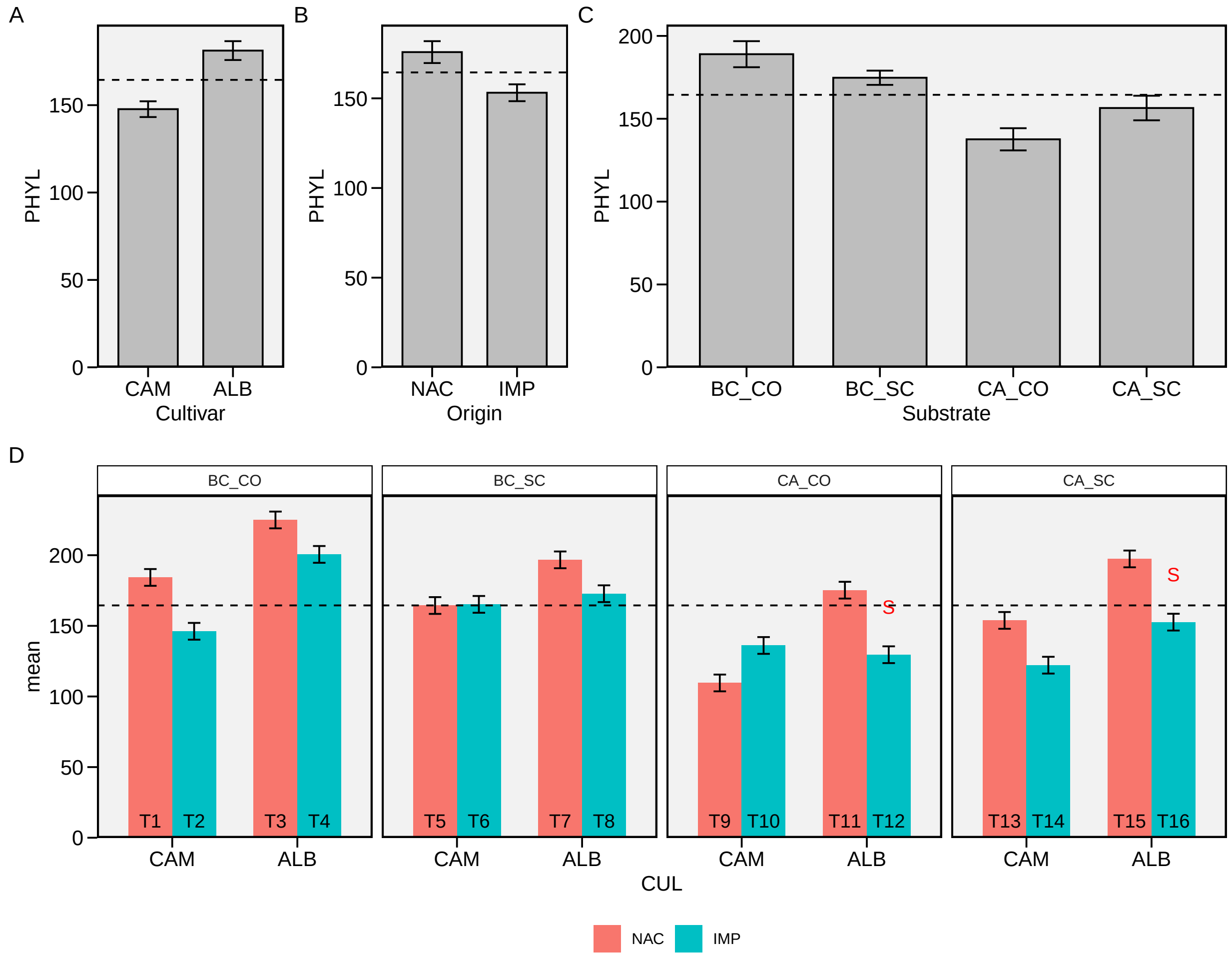

Figure S 8: Phyllochron. Main effect of cultivar (A), origin (B), substrate (C) and for the three-way interaction (D). The horizonal dashed line shows the overall mean. The 'S' letter indicates the selected treatments. Bars shows the mean and standard error. N = 4.

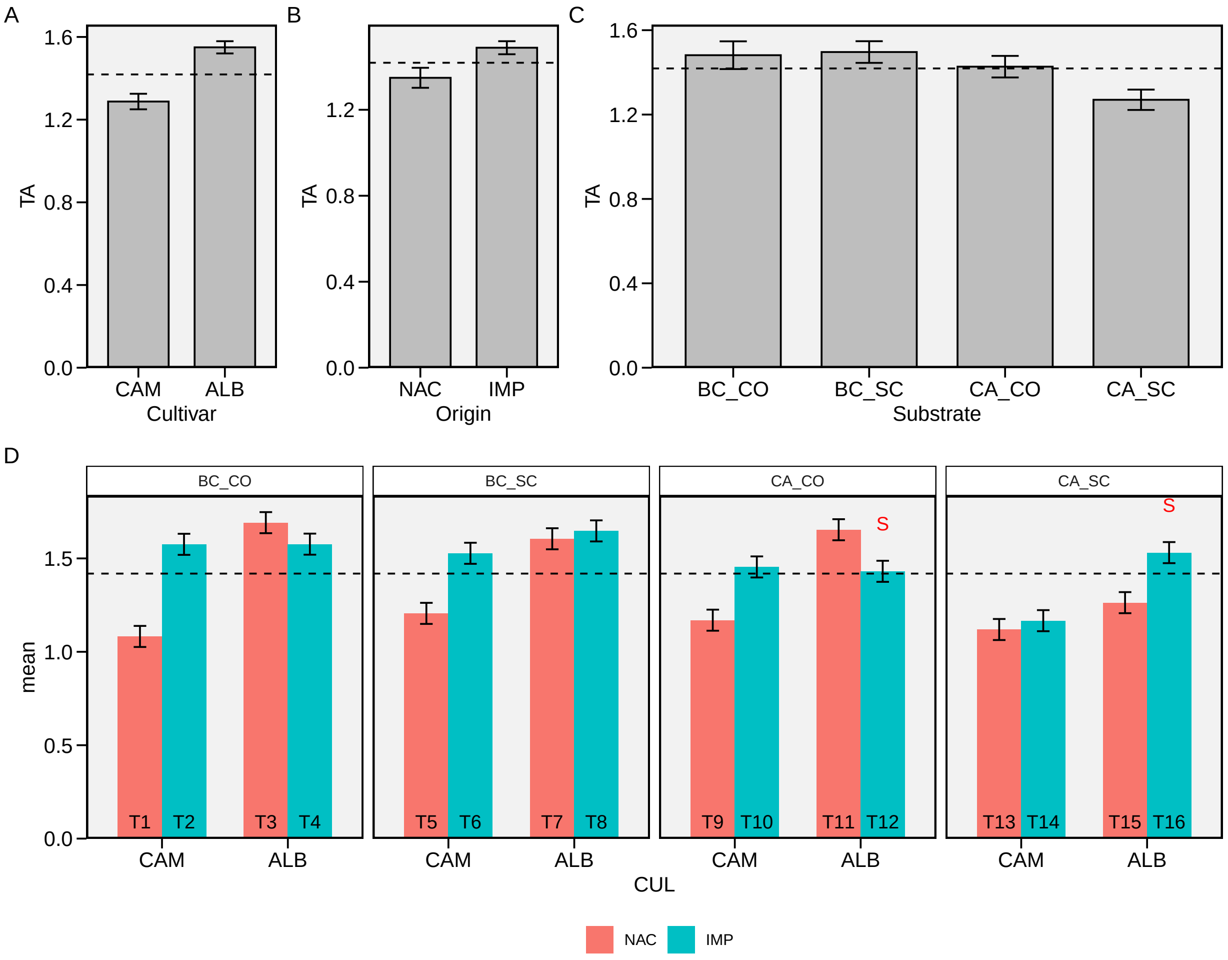

Figure S 9: Total acid. Main effect of cultivar (A), origin (B), substrate (C) and for the three-way interaction (D). The horizonal dashed line shows the overall mean. The ’S’ letter indicates the selected treatments. Bars shows the mean and standard error. N = 4.

#### 2.3 Traits with positive desired gains

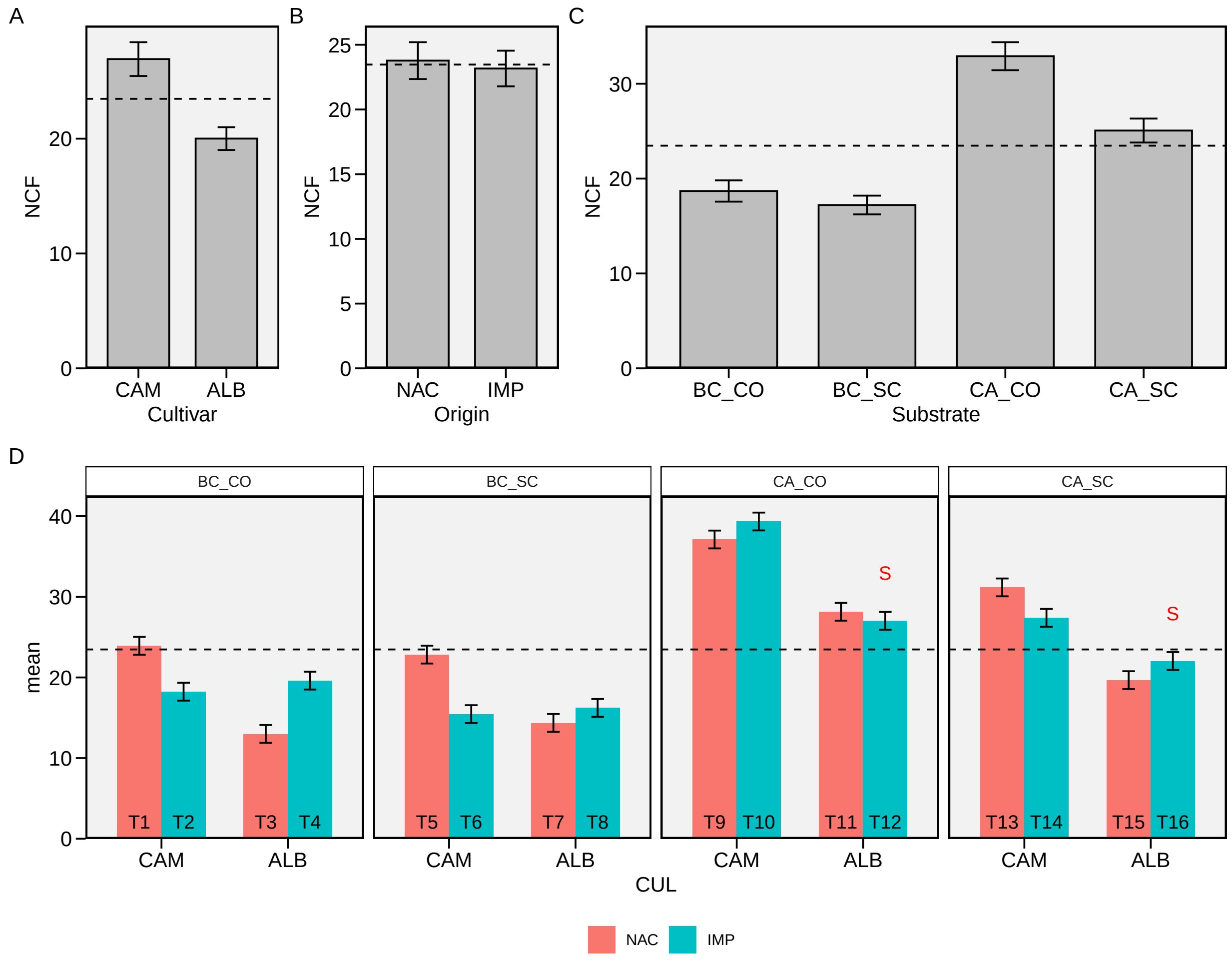

Figure S 10: Number of commercial fruits. Main effect of cultivar (A), origin (B), substrate (C) and for the three-way interaction (D). The horizonal dashed line shows the overall mean. The 'S' letter indicates the selected treatments. Bars shows the mean and standard error. N = 4.

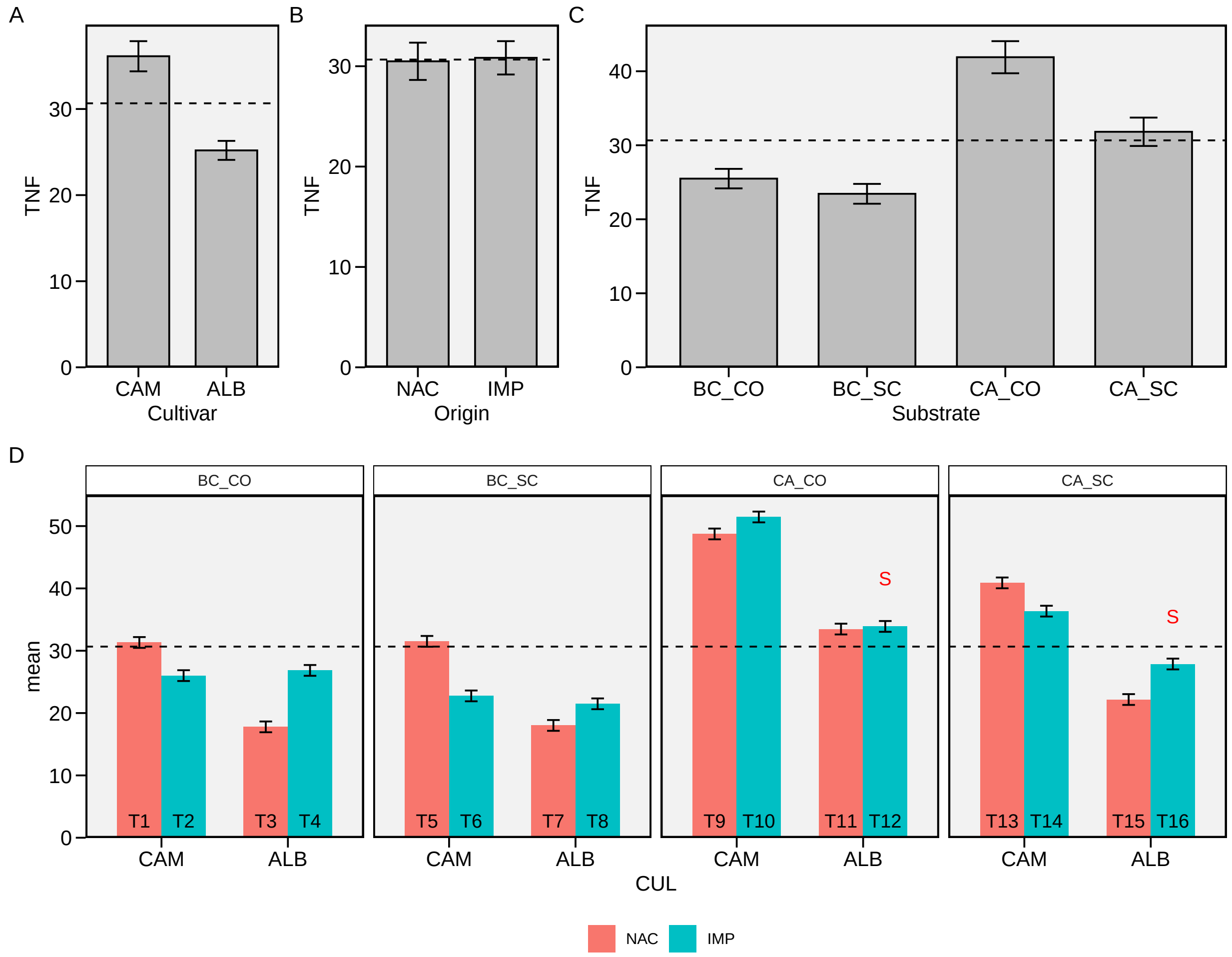

Figure S 11: Total number of fruits. Main effect of cultivar (A), origin (B), substrate (C) and for the three-way interaction (D). The horizonal dashed line shows the overall mean. The 'S' letter indicates the selected treatments. Bars shows the mean and standard error. N = 4.

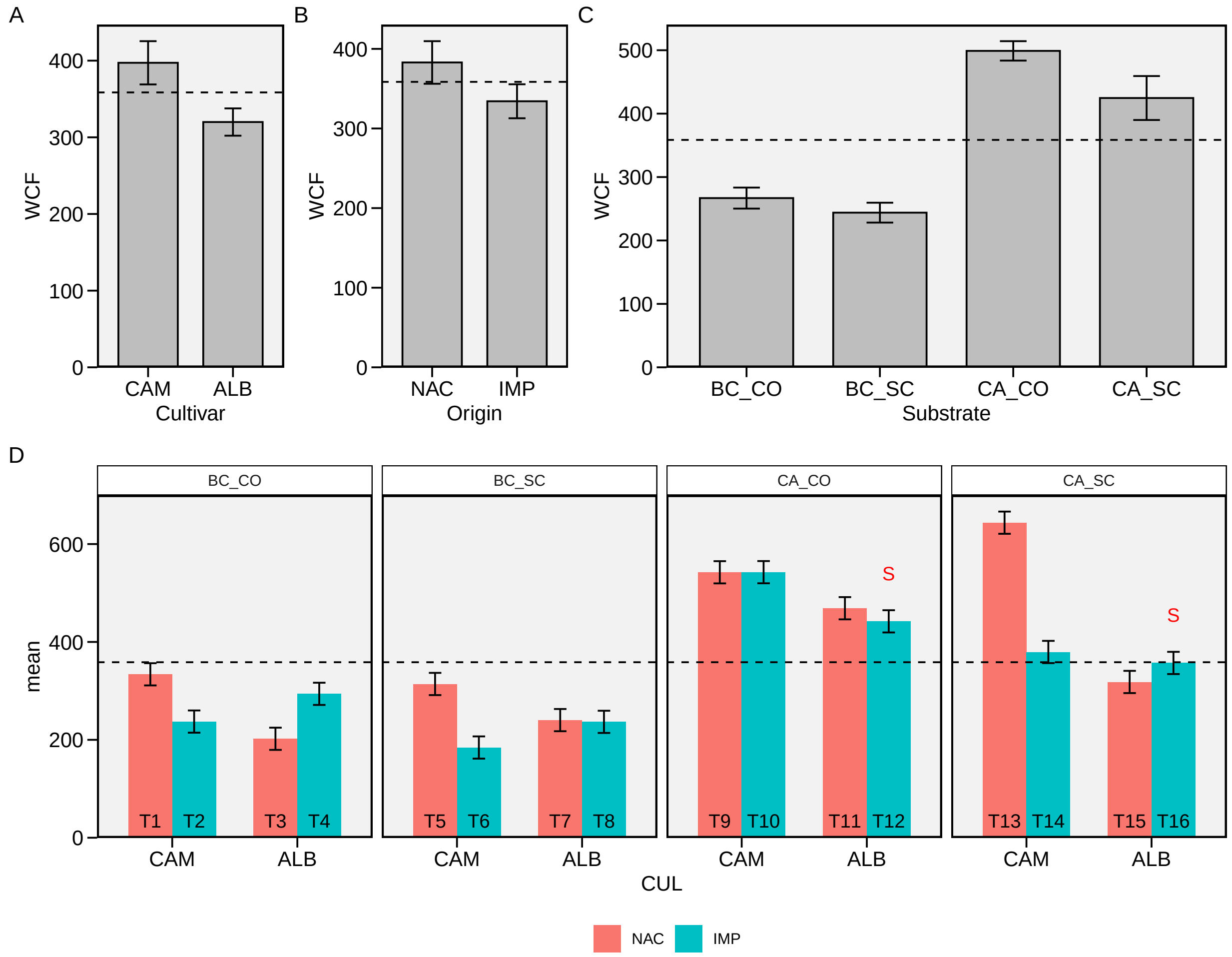

Figure S 12: Weight of commercial fruits. Main effect of cultivar (A), origin (B), substrate (C) and for the three-way interaction (D). The horizonal dashed line shows the overall mean. The 'S' letter indicates the selected treatments. Bars shows the mean and standard error. N = 4.

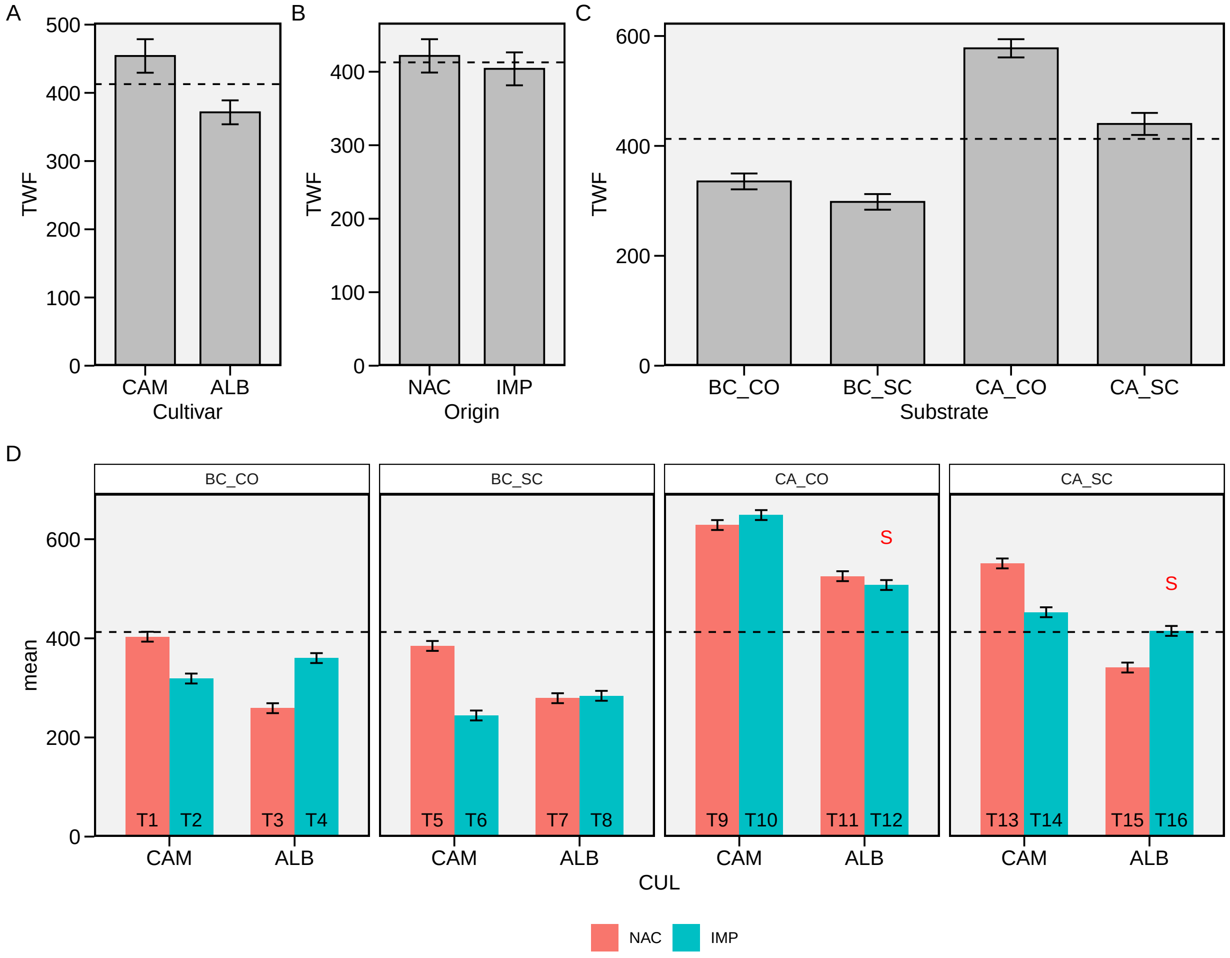

Figure S 13: Total weight of fruits. Main effect of cultivar (A), origin (B), substrate (C) and for the three-way interaction (D). The horizonal dashed line shows the overall mean. The 'S' letter indicates the selected treatments. Bars shows the mean and standard error. N = 4.

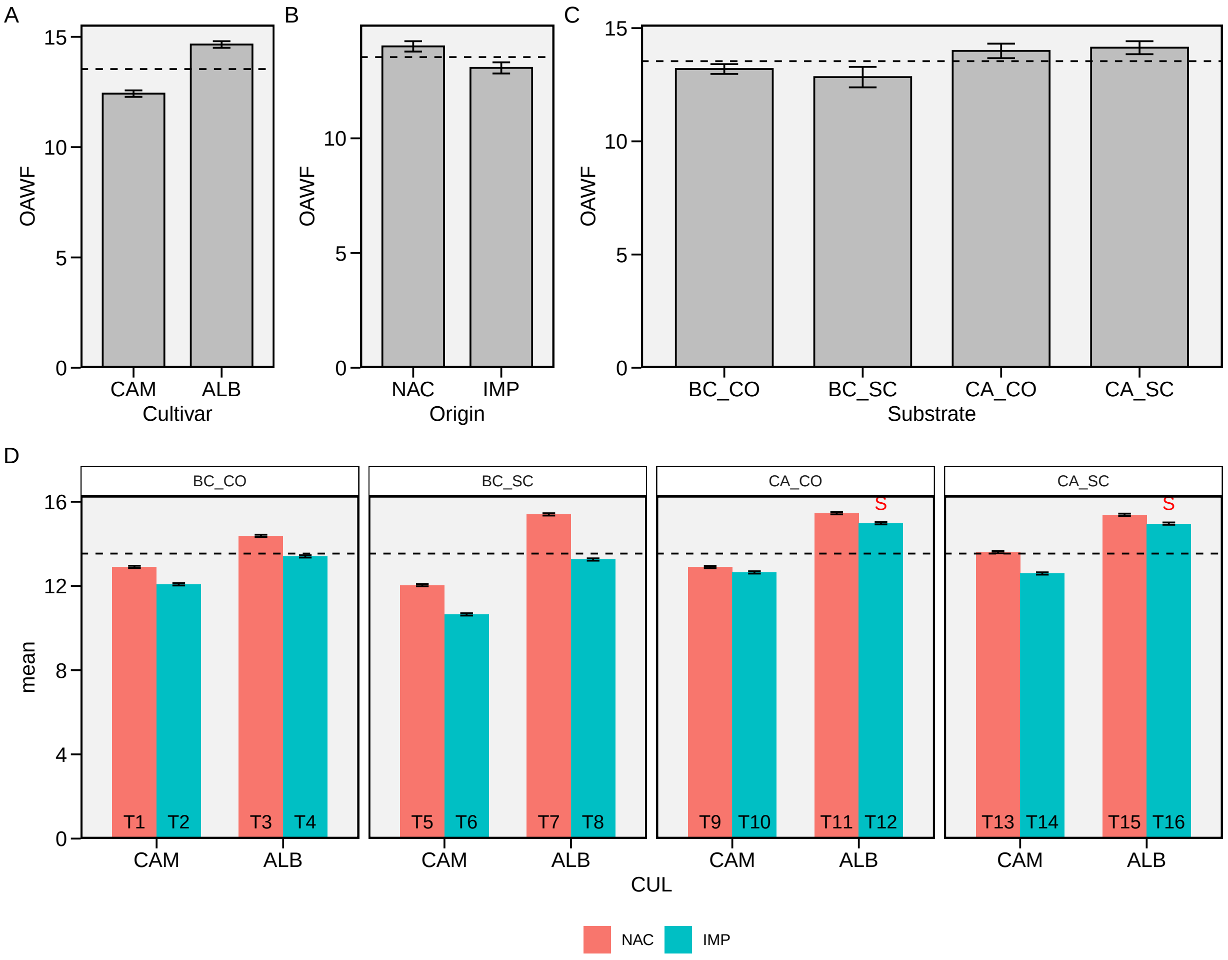

Figure S 14: Overall average of weigth of fruits. Main effect of cultivar (A), origin (B), substrate (C) and for the three-way interaction (D). The horizonal dashed line shows the overall mean. The 'S' letter indicates the selected treatments. Bars shows the mean and standard error. N = 4.

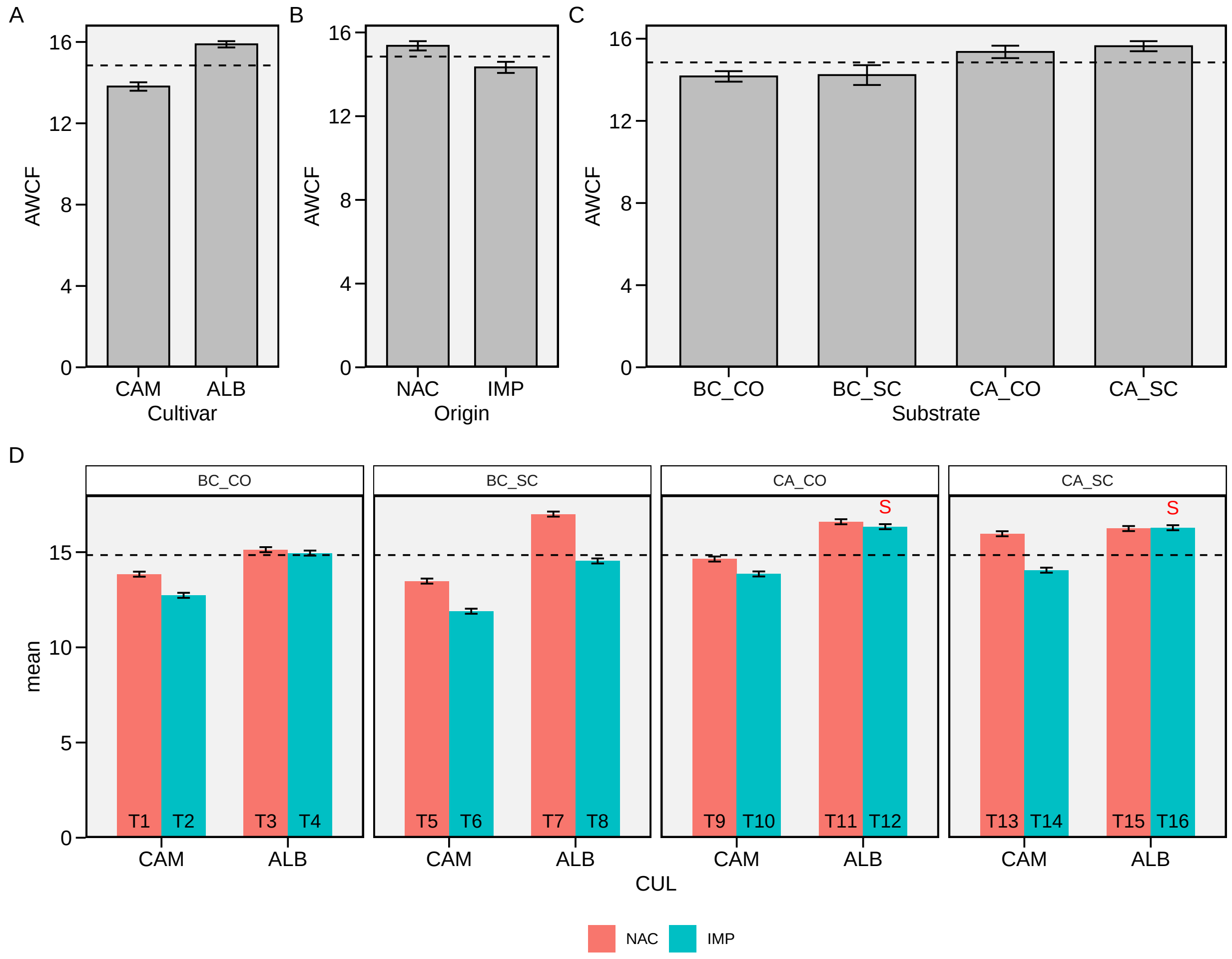

Figure S 15: Average weigth of comercial fruits. Main effect of cultivar (A), origin (B), substrate (C) and for the three-way interaction (D). The horizonal dashed line shows the overall mean. The ’S’ letter indicates the selected treatments. Bars shows the mean and standard error. N = 4.

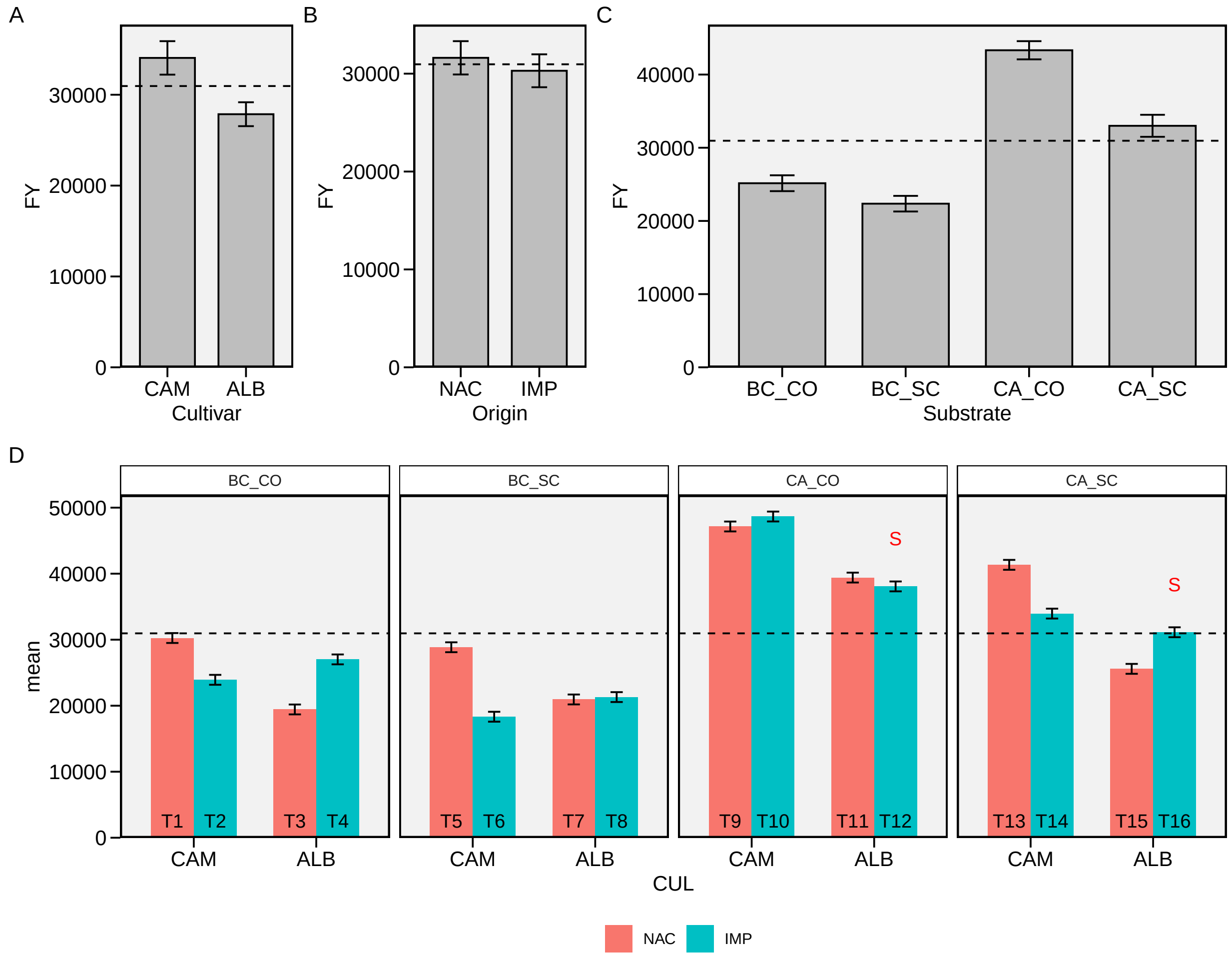

Figure S 16: Fruit yield. Main effect of cultivar (A), origin (B), substrate (C) and for the three-way interaction (D). The horizonal dashed line shows the overall mean. The 'S' letter indicates the selected treatments. Bars shows the mean and standard error. N = 4.

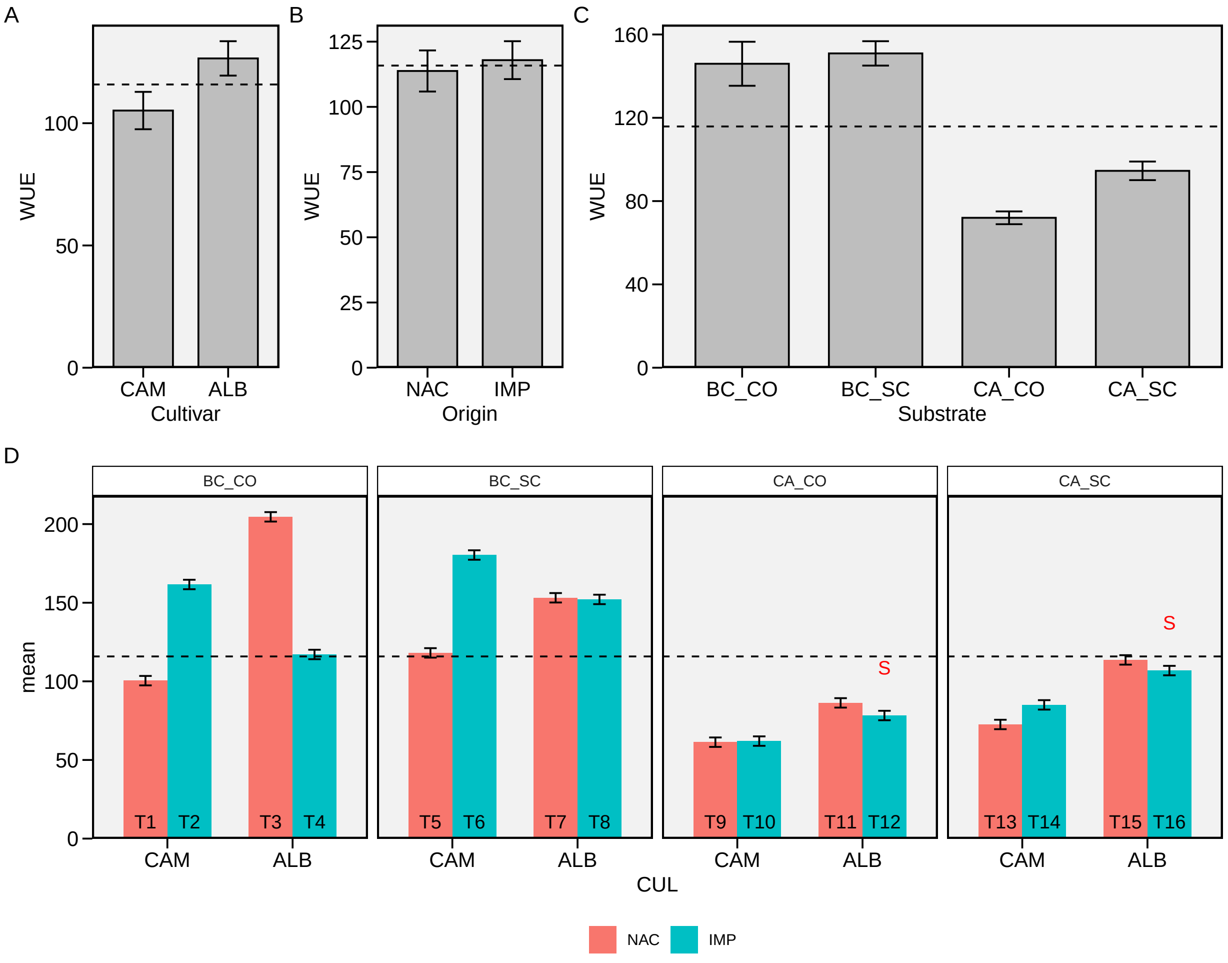

Figure S 17: Water use efficiency. Main effect of cultivar (A), origin (B), substrate (C) and for the three-way interaction (D). The horizonal dashed line shows the overall mean. The 'S' letter indicates the selected treatments. Bars shows the mean and standard error. N = 4.

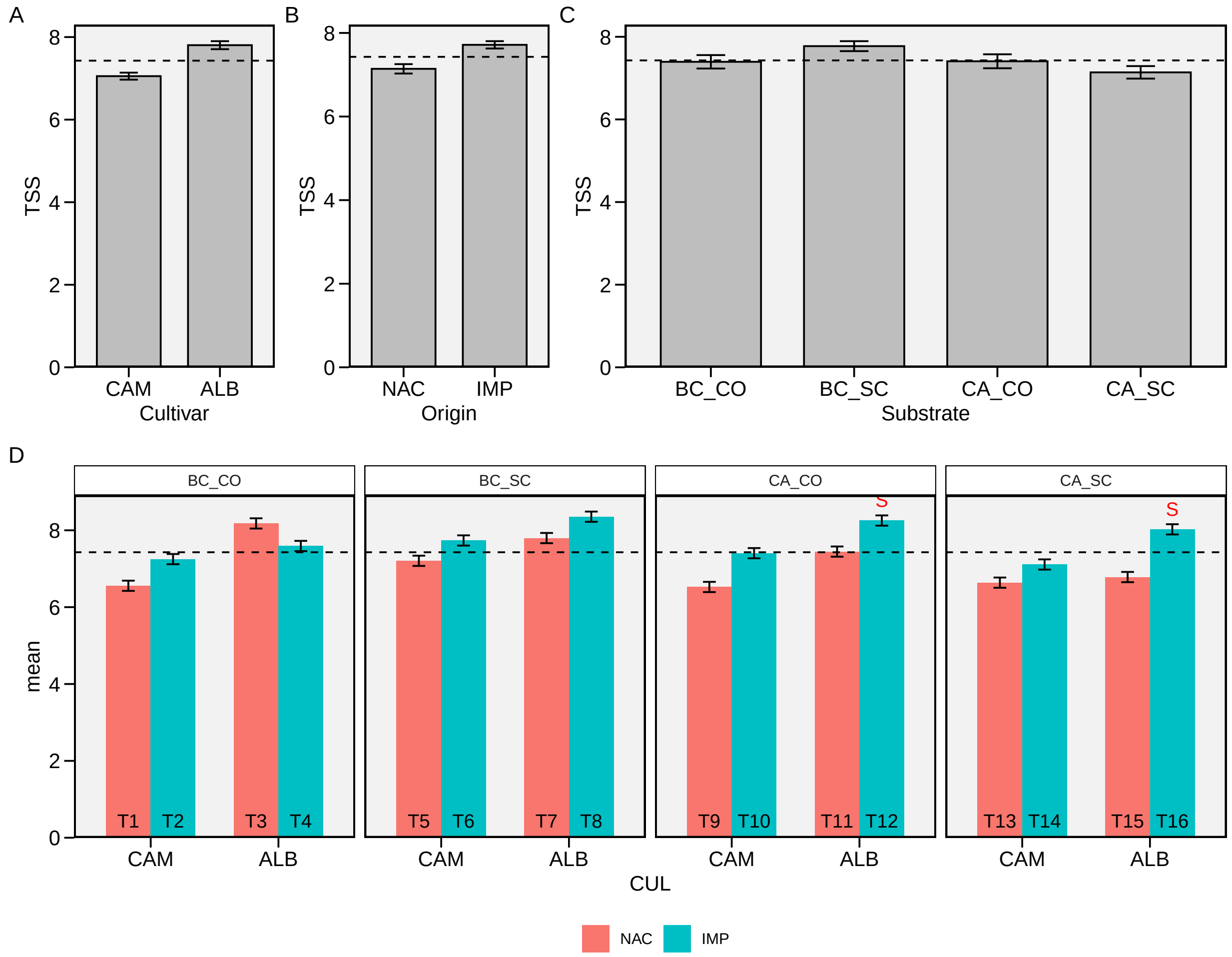

Figure S 18: Total soluble solids. Main effect of cultivar (A), origin (B), substrate (C) and for the three-way interaction (D). The horizonal dashed line shows the overall mean. The ’S’ letter indicates the selected treatments. Bars shows the mean and standard error. N = 4.

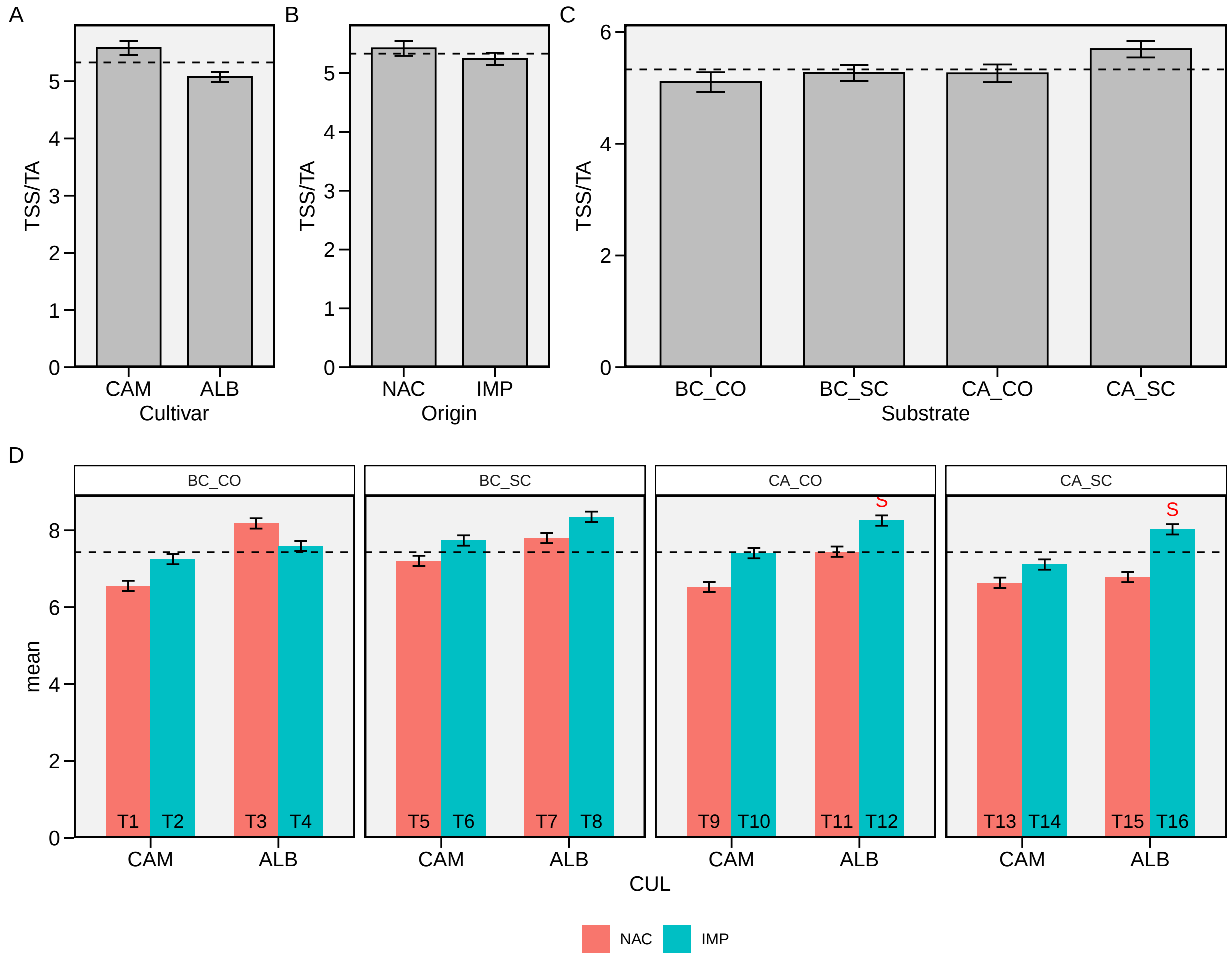

Figure S 19: TSS/TA ratio. Main effect of cultivar (A), origin (B), substrate (C) and for the three-way interaction (D). The horizonal dashed line shows the overall mean. The 'S' letter indicates the selected treatments. Bars shows the mean and standard error. N = 4.

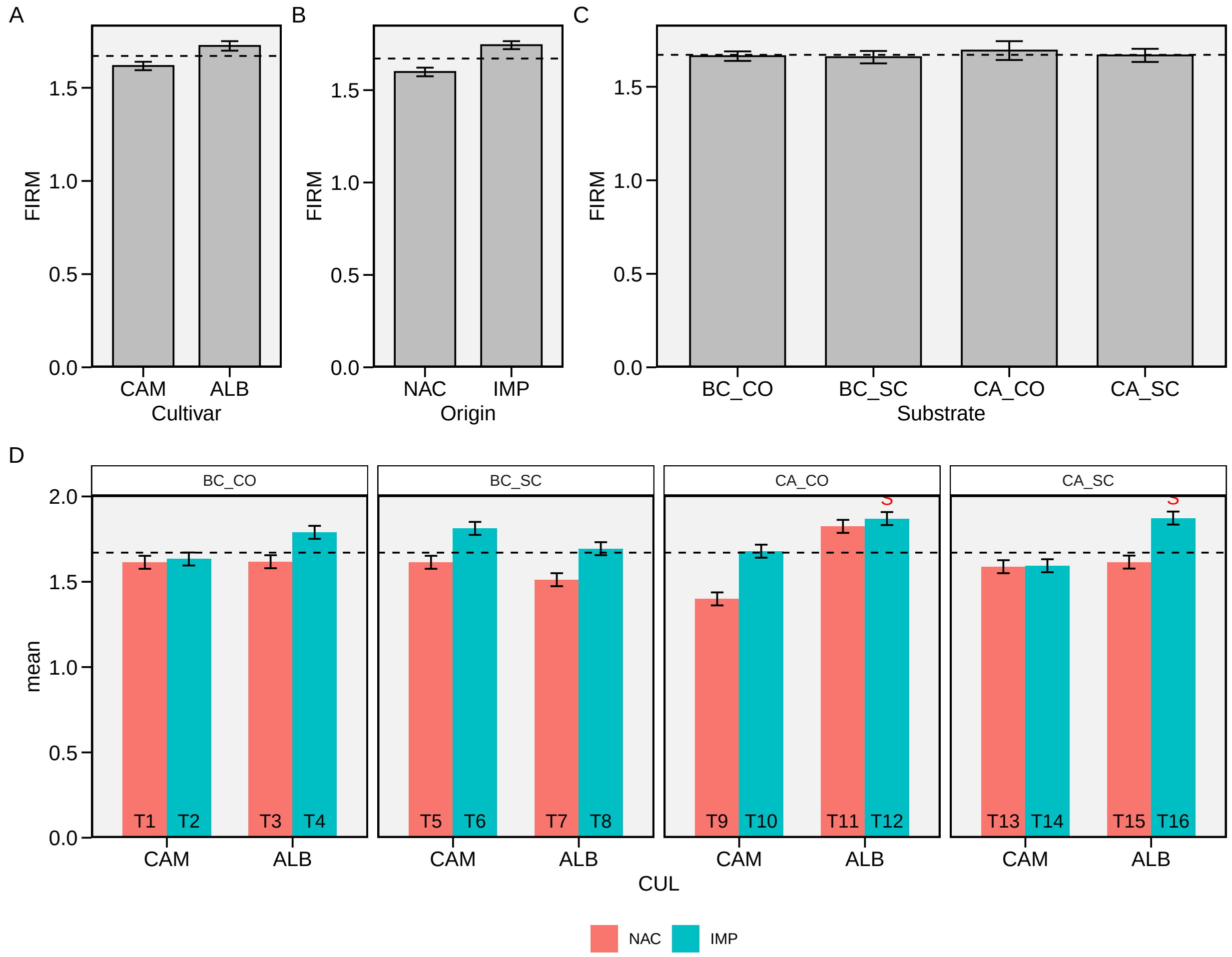

Figure S 20: Flesh firmness. Main effect of cultivar (A), origin (B), substrate (C) and for the three-way interaction (D). The horizonal dashed line shows the overall mean. The ’S’ letter indicates the selected treatments. Bars shows the mean and standard error. N = 4.

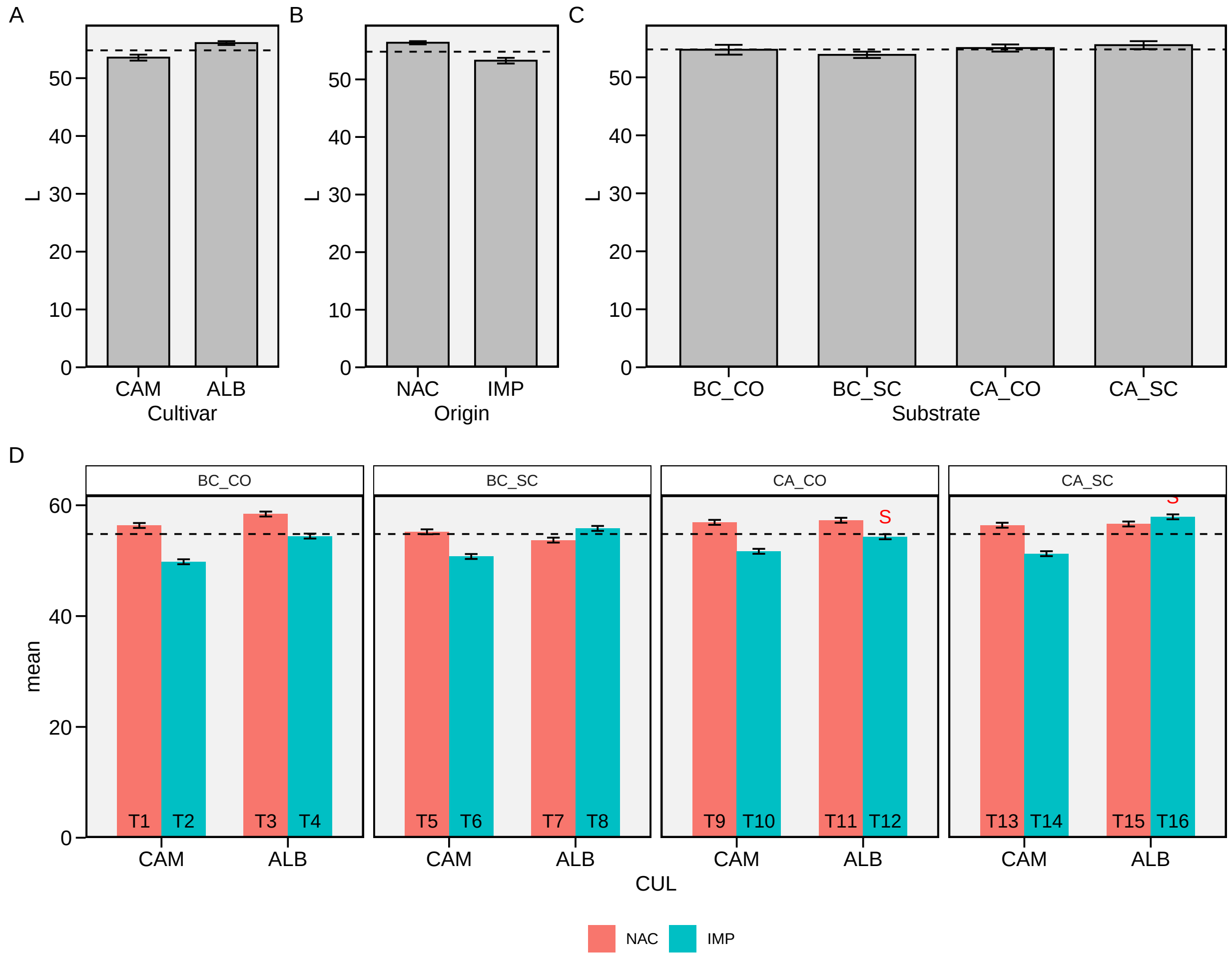

Figure S 21: L. Main effect of cultivar (A), origin (B), substrate (C) and for the three-way interaction (D). The horizonal dashed line shows the overall mean. The 'S' letter indicates the selected treatments. Bars shows the mean and standard error. N = 4.

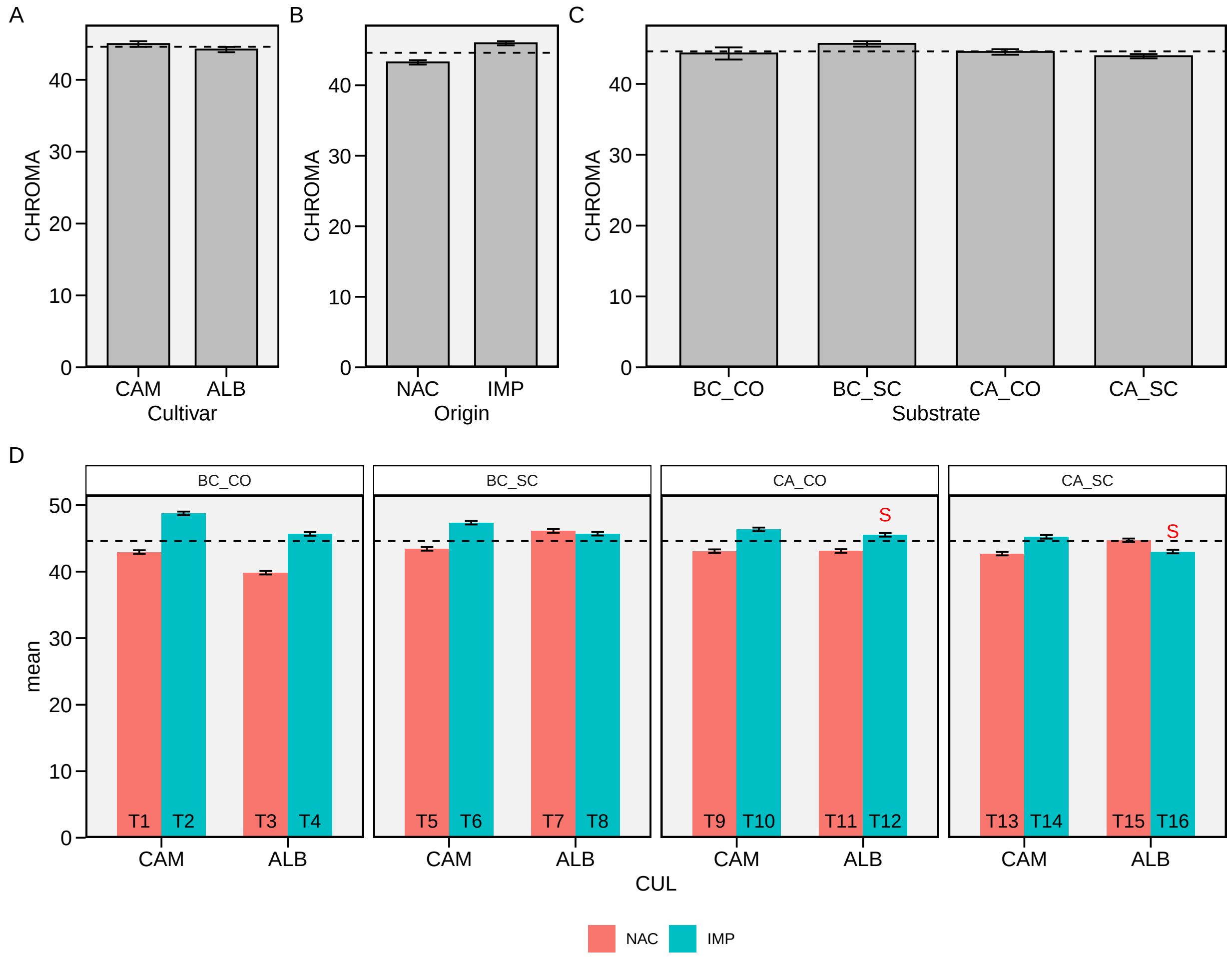

Figure S 22: CHROMA. Main effect of cultivar (A), origin (B), substrate (C) and for the three-way interaction (D). The horizonal dashed line shows the overall mean. The 'S' letter indicates the selected treatments. Bars shows the mean and standard error. N = 4.

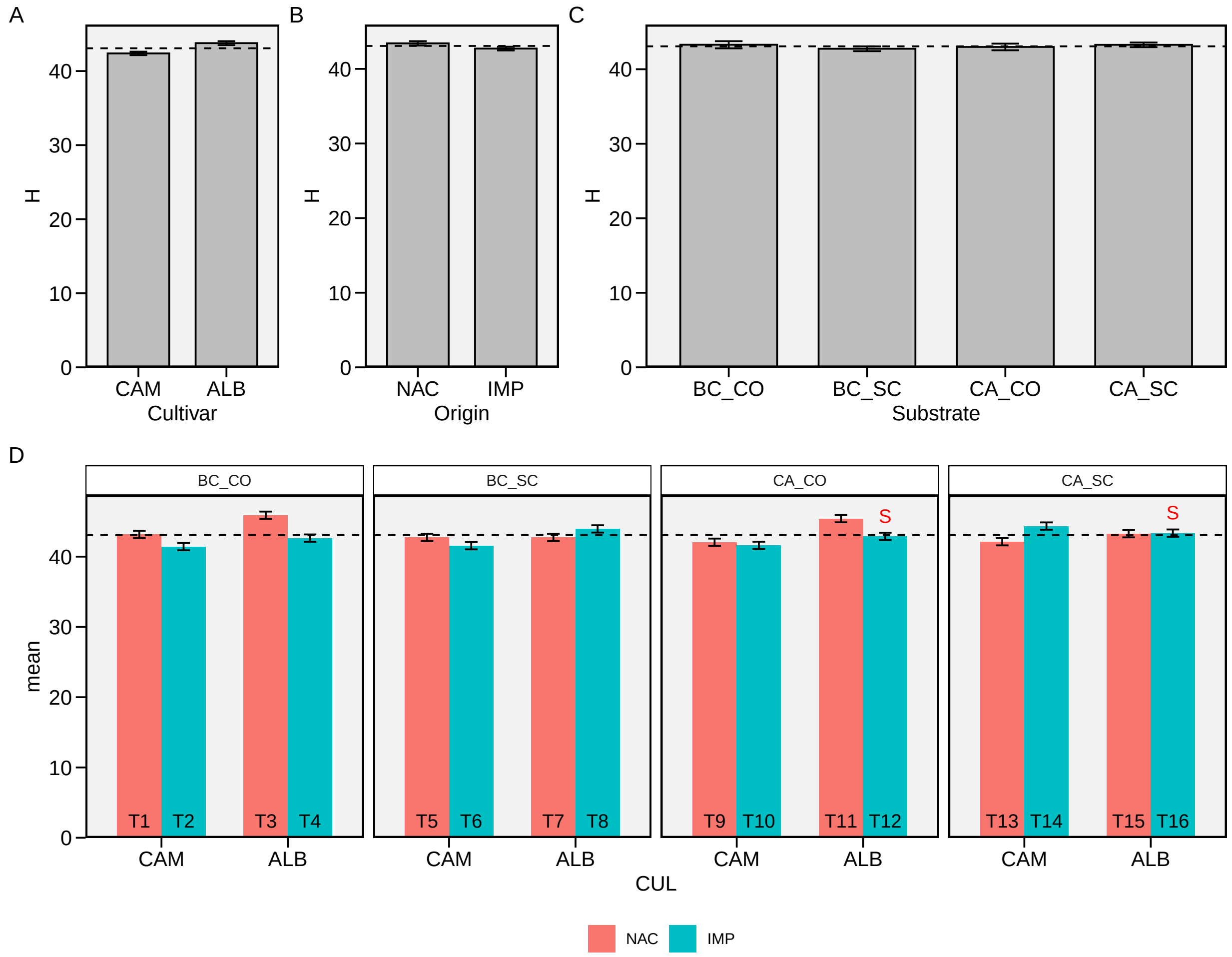

Figure S 23: H Main effect of cultivar (A), origin (B), substrate (C) and for the three-way interaction (D). The horizonal dashed line shows the overall mean. The 'S' letter indicates the selected treatments. Bars shows the mean and standard error. N = 4.

### 3 Appendix C - Supplementary tables

Supplementary Table S 1: Doses of mineral fertilizers in 1000 liters of water for strawberry fertigation in a substrate of pH 7 or above.

| Fertilizers | Vegetative Phase | Reproductive Phase |
|---|---|---|
| Calcium nitrate (15.5-00-00) | 480 g | 480 g |
| Potassium nitrate (12-00-45) | 300 g | 180 g |
| Magnesium Sulphate (00-00-00-09) | 360 g | 360 g |
| Ammonium sulfate (20-00-00) | 50 g | 70 g |
| Potassium Sulphate (00-00-50) | 70 g | 260 g |
| Phosphoric acid (85%) | 110 ml | 110 ml |
| Boric acid(17%B) | 1.8 g | 1.8 g |
| Copper sulphate (25%Cu) | 0.18 g | 0.18 g |
| Manganese Sulfate (25%Mn) | 1.2 g | 1.2 g |
| Zinc sulfate (20% Zn) | 0.6 g | 0.6 g |
| Sodium Molybdate (39%Mo) | 0.18 g | 0.18 g |
| Chelated iron (6% Fe) | 36 g | 36 g |

Supplementary Table S 2: Density (DS), total porosity (TP), aeration space (AS), readily available water (RAW), buffer water (BW), available water (AW) and remaining water (AR) for the substrates used: S1: Sugarcane bagasse + organic compost; S2: Sugarcane bagasse + commercial substrate - Carolina; S3: Rice husk + organic compost; and S4: Rice husk + commercial substrate - Carolina.

| Substracts | DS | TP | AS | RAW | BW | AW | RW |
|---|---|---|---|---|---|---|---|
| S1 | 0.14 | 0.74 | 0.48 | 0.05 | 0.01 | 0.06 | 0.20 |
| S2 | 0.09 | 0.75 | 0.46 | 0.09 | 0.02 | 0.11 | 0.18 |
| S3 | 0.23 | 0.63 | 0.27 | 0.15 | 0.03 | 0.18 | 0.18 |
| S4 | 0.13 | 0.67 | 0.29 | 0.24 | 0.08 | 0.32 | 0.05 |

Supplementary Table S 3: Analysis of variance table for 23 strawberry traits showing the significance for the effects of cultivar (CUL), origin (ORI), substrate (SUB), and its respective two- and three-way interactions. Bold values indicates significant (p-value < 0.05) effects.

| term | TRAIT | df | statistic | p.value |
|---|---|---|---|---|
| **CUL** | **NNCF** | **1** | **50.359172** | **0.000000** |
| **CUL** | **WNCF** | **1** | **17.429620** | **0.000135** |
| CUL | AWNCF | 1 | 1.715339 | 0.196941 |
| **CUL** | **WUE** | **1** | **4.142216** | **0.047742** |
| **CUL** | **NDBF** | **1** | **25.168011** | **0.000009** |
| **CUL** | **NDFF** | **1** | **51.995429** | **0.000000** |
| **CUL** | **NDBH** | **1** | **27.097689** | **0.000005** |
| **CUL** | **PHYL** | **1** | **11.208413** | **0.001653** |
| **CUL** | **TA** | **1** | **31.922404** | **0.000001** |
| **CUL** | **NCF** | **1** | **18.275987** | **0.000098** |
| **CUL** | **TNF** | **1** | **37.063715** | **0.000000** |
| **CUL** | **WCF** | **1** | **5.734357** | **0.020862** |
| **CUL** | **TWF** | **1** | **10.644977** | **0.002110** |
| **CUL** | **AWCF** | **1** | **37.954836** | **0.000000** |
| **CUL** | **OAWF** | **1** | **55.404437** | **0.000000** |
| **CUL** | **FY** | **1** | **10.644977** | **0.002110** |
| **CUL** | **TSS** | **1** | **16.059800** | **0.000228** |
| **CUL** | **TSS/TA** | **1** | **8.269576** | **0.006139** |
| CUL | FIRM | 1 | 2.102635 | 0.153980 |
| **CUL** | **L** | **1** | **21.842798** | **0.000027** |
| CUL | CHROMA | 1 | 0.831076 | 0.366820 |
| CUL | H | 1 | 2.404527 | 0.127990 |
| ORI | NNCF | 1 | 3.036952 | 0.088219 |
| ORI | WNCF | 1 | 2.027685 | 0.161353 |
| ORI | AWNCF | 1 | 0.069242 | 0.793645 |
| ORI | WUE | 1 | 0.156647 | 0.694133 |
| **ORI** | **NDBF** | **1** | **58.094492** | **0.000000** |
| **ORI** | **NDFF** | **1** | **31.257361** | **0.000001** |
| **ORI** | **NDBH** | **1** | **56.537154** | **0.000000** |
| **ORI** | **PHYL** | **1** | **5.125225** | **0.028449** |
| **ORI** | **TA** | **1** | **9.089844** | **0.004214** |
| ORI | NCF | 1 | 0.141948 | 0.708122 |
| ORI | TNF | 1 | 0.037212 | 0.847902 |
| ORI | WCF | 1 | 2.281562 | 0.137911 |
| ORI | TWF | 1 | 0.485235 | 0.489643 |
| **ORI** | **AWCF** | **1** | **9.358983** | **0.003731** |
| **ORI** | **OAWF** | **1** | **9.824985** | **0.003029** |
| ORI | FY | 1 | 0.485235 | 0.489643 |
| **ORI** | **TSS** | **1** | **9.394378** | **0.003672** |

Supplementary Table S 3: Analysis of variance table for 23 strawberry traits showing the significance for the effects of cultivar (CUL), origin (ORI), substrate (SUB), and its respective two- and three-way interactions. Bold values indicates significant (p-value < 0.05) effects. *(continued)*

| term | TRAIT | df | statistic | p.value |
|---|---|---|---|---|
| ORI | TSS/TA | 1 | 0.906289 | 0.346187 |
| ORI | FIRM | 1 | 3.821482 | 0.056828 |
| **ORI** | **L** | **1** | **33.449767** | **0.000001** |
| **ORI** | **CHROMA** | **1** | **10.266933** | **0.002490** |
| ORI | H | 1 | 0.615373 | 0.436882 |
| **ORI:CUL** | **NNCF** | **1** | **5.316668** | **0.025786** |
| ORI:CUL | WNCF | 1 | 1.029794 | 0.315632 |
| ORI:CUL | AWNCF | 1 | 0.015272 | 0.902197 |
| **ORI:CUL** | **WUE** | **1** | **8.191849** | **0.006365** |
| **ORI:CUL** | **NDBF** | **1** | **9.489324** | **0.003519** |
| **ORI:CUL** | **NDFF** | **1** | **12.786056** | **0.000848** |
| **ORI:CUL** | **NDBH** | **1** | **6.107417** | **0.017312** |
| ORI:CUL | PHYL | 1 | 1.446386 | 0.235399 |
| **ORI:CUL** | **TA** | **1** | **10.012846** | **0.002786** |
| ORI:CUL | NCF | 1 | 3.526766 | 0.066875 |
| **ORI:CUL** | **TNF** | **1** | **5.772567** | **0.020465** |
| **ORI:CUL** | **WCF** | **1** | **5.237072** | **0.026859** |
| **ORI:CUL** | **TWF** | **1** | **5.270793** | **0.026399** |
| ORI:CUL | AWCF | 1 | 0.882967 | 0.352407 |
| ORI:CUL | OAWF | 1 | 0.050038 | 0.824009 |
| **ORI:CUL** | **FY** | **1** | **5.270793** | **0.026399** |
| ORI:CUL | TSS | 1 | 0.137744 | 0.712276 |
| **ORI:CUL** | **TSS/TA** | **1** | **7.667970** | **0.008136** |
| ORI:CUL | FIRM | 1 | 0.067509 | 0.796183 |
| **ORI:CUL** | **L** | **1** | **17.057719** | **0.000155** |
| ORI:CUL | CHROMA | 1 | 1.924919 | 0.172148 |
| ORI:CUL | H | 1 | 0.217943 | 0.642865 |
| **SUB** | **NNCF** | **3** | **4.721638** | **0.005998** |
| SUB | WNCF | 3 | 2.668227 | 0.058928 |
| SUB | AWNCF | 3 | 1.841424 | 0.153231 |
| **SUB** | **WUE** | **3** | **13.703015** | **0.000002** |
| **SUB** | **NDBF** | **3** | **8.915885** | **0.000095** |
| **SUB** | **NDFF** | **3** | **11.401853** | **0.000011** |
| SUB | NDBH | 3 | 2.384988 | 0.081657 |
| **SUB** | **PHYL** | **3** | **4.962146** | **0.004640** |
| **SUB** | **TA** | **3** | **4.968764** | **0.004607** |
| **SUB** | **NCF** | **3** | **19.504522** | **0.000000** |
| **SUB** | **TNF** | **3** | **21.328678** | **0.000000** |

Supplementary Table S 3: Analysis of variance table for 23 strawberry traits showing the significance for the effects of cultivar (CUL), origin (ORI), substrate (SUB), and its respective two- and three-way interactions. Bold values indicates significant (p-value < 0.05) effects. *(continued)*

| term | TRAIT | df | statistic | p.value |
|---|---|---|---|---|
| **SUB** | **WCF** | **3** | **14.610909** | **0.000001** |
| **SUB** | **TWF** | **3** | **24.461840** | **0.000000** |
| **SUB** | **AWCF** | **3** | **5.079991** | **0.004095** |
| **SUB** | **OAWF** | **3** | **4.387025** | **0.008606** |
| **SUB** | **FY** | **3** | **24.461840** | **0.000000** |
| SUB | TSS | 3 | 1.940997 | 0.136527 |
| SUB | TSS/TA | 3 | 1.541182 | 0.216868 |
| SUB | FIRM | 3 | 0.044157 | 0.987503 |
| SUB | L | 3 | 1.702448 | 0.179993 |
| SUB | CHROMA | 3 | 0.772610 | 0.515422 |
| SUB | H | 3 | 0.082386 | 0.969282 |
| SUB:CUL | NNCF | 3 | 2.759285 | 0.053084 |
| SUB:CUL | WNCF | 3 | 0.720274 | 0.545127 |
| SUB:CUL | AWNCF | 3 | 0.134507 | 0.938971 |
| SUB:CUL | WUE | 3 | 0.376755 | 0.770183 |
| SUB:CUL | NDBF | 3 | 1.472743 | 0.234665 |
| **SUB:CUL** | **NDFF** | **3** | **4.273727** | **0.009736** |
| SUB:CUL | NDBH | 3 | 0.814589 | 0.492556 |
| SUB:CUL | PHYL | 3 | 0.340520 | 0.796114 |
| SUB:CUL | TA | 3 | 0.113060 | 0.952019 |
| SUB:CUL | NCF | 3 | 0.953504 | 0.422900 |
| SUB:CUL | TNF | 3 | 1.826151 | 0.155967 |
| SUB:CUL | WCF | 3 | 1.236264 | 0.307667 |
| SUB:CUL | TWF | 3 | 0.872067 | 0.462625 |
| SUB:CUL | AWCF | 3 | 1.329663 | 0.276556 |
| SUB:CUL | OAWF | 3 | 1.236531 | 0.307573 |
| SUB:CUL | FY | 3 | 0.872067 | 0.462625 |
| SUB:CUL | TSS | 3 | 0.336121 | 0.799270 |
| SUB:CUL | TSS/TA | 3 | 0.400071 | 0.753604 |
| SUB:CUL | FIRM | 3 | 1.369146 | 0.264327 |
| SUB:CUL | L | 3 | 0.892545 | 0.452343 |
| SUB:CUL | CHROMA | 3 | 0.884308 | 0.456454 |
| SUB:CUL | H | 3 | 0.309953 | 0.818063 |
| SUB:ORI | NNCF | 3 | 0.286130 | 0.835152 |
| SUB:ORI | WNCF | 3 | 0.294717 | 0.828998 |
| SUB:ORI | AWNCF | 3 | 0.038869 | 0.989627 |
| SUB:ORI | WUE | 3 | 0.806417 | 0.496940 |
| SUB:ORI | NDBF | 3 | 2.098238 | 0.113781 |

Supplementary Table S 3: Analysis of variance table for 23 strawberry traits showing the significance for the effects of cultivar (CUL), origin (ORI), substrate (SUB), and its respective two- and three-way interactions. Bold values indicates significant (p-value < 0.05) effects. *(continued)*

| term | TRAIT | df | statistic | p.value |
|---|---|---|---|---|
| SUB:ORI | NDFF | 3 | 0.766385 | 0.518885 |
| SUB:ORI | NDBH | 3 | 0.433662 | 0.729936 |
| SUB:ORI | PHYL | 3 | 0.507570 | 0.679054 |
| SUB:ORI | TA | 3 | 0.630449 | 0.599178 |
| SUB:ORI | NCF | 3 | 0.226592 | 0.877403 |
| SUB:ORI | TNF | 3 | 0.331348 | 0.802697 |
| SUB:ORI | WCF | 3 | 0.625393 | 0.602333 |
| SUB:ORI | TWF | 3 | 0.463800 | 0.708967 |
| SUB:ORI | AWCF | 3 | 1.027539 | 0.389431 |
| SUB:ORI | OAWF | 3 | 0.988191 | 0.406912 |
| SUB:ORI | FY | 3 | 0.463800 | 0.708967 |
| SUB:ORI | TSS | 3 | 1.009803 | 0.397225 |
| SUB:ORI | TSS/TA | 3 | 1.836398 | 0.154126 |
| SUB:ORI | FIRM | 3 | 0.074938 | 0.973164 |
| **SUB:ORI** | **L** | **3** | **3.098406** | **0.036053** |
| SUB:ORI | CHROMA | 3 | 1.838414 | 0.153766 |
| SUB:ORI | H | 3 | 0.810766 | 0.494602 |
| SUB:ORI:CUL | NNCF | 3 | 0.370192 | 0.774866 |
| SUB:ORI:CUL | WNCF | 3 | 1.025515 | 0.390314 |
| SUB:ORI:CUL | AWNCF | 3 | 0.228099 | 0.876348 |
| SUB:ORI:CUL | WUE | 3 | 2.307750 | 0.089279 |
| SUB:ORI:CUL | NDBF | 3 | 0.643983 | 0.590790 |
| SUB:ORI:CUL | NDFF | 3 | 0.416692 | 0.741858 |
| SUB:ORI:CUL | NDBH | 3 | 0.312421 | 0.816290 |
| SUB:ORI:CUL | PHYL | 3 | 0.804888 | 0.497764 |
| **SUB:ORI:CUL** | **TA** | **3** | **3.956537** | **0.013789** |
| SUB:ORI:CUL | NCF | 3 | 1.094160 | 0.361379 |
| SUB:ORI:CUL | TNF | 3 | 1.084226 | 0.365442 |
| SUB:ORI:CUL | WCF | 3 | 1.135546 | 0.344899 |
| SUB:ORI:CUL | TWF | 3 | 1.058504 | 0.376155 |
| SUB:ORI:CUL | AWCF | 3 | 0.759336 | 0.522830 |
| SUB:ORI:CUL | OAWF | 3 | 0.212597 | 0.887153 |
| SUB:ORI:CUL | FY | 3 | 1.058504 | 0.376155 |
| SUB:ORI:CUL | TSS | 3 | 1.282691 | 0.291808 |
| SUB:ORI:CUL | TSS/TA | 3 | 1.975013 | 0.131249 |
| SUB:ORI:CUL | FIRM | 3 | 0.514752 | 0.674212 |
| SUB:ORI:CUL | L | 3 | 1.204667 | 0.318921 |
| SUB:ORI:CUL | CHROMA | 3 | 0.443449 | 0.723096 |

Supplementary Table S 3: Analysis of variance table for 23 strawberry traits showing the significance for the effects of cultivar (CUL), origin (ORI), substrate (SUB), and its respective two- and three-way interactions. Bold values indicates significant (p-value $< 0.05$) effects. *(continued)*

| term | TRAIT | df | statistic | p.value |
|---|---|---|---|---|
| SUB:ORI:CUL | H | 3 | 0.366726 | 0.777343 |

Supplementary Table S 4: Eigenvalues, explained variance, and cummulated variance for the factor analysis perfomed with 23 strawberry traits.

| Factor | Eigenvalue | Explained variance (%) | Cummulative variance (%) |
|---|---|---|---|
| PC1 | 8.752 | 39.782 | 39.782 |
| PC2 | 4.726 | 21.481 | 61.263 |
| PC3 | 3.680 | 16.727 | 77.990 |
| PC4 | 1.501 | 6.822 | 84.813 |
| PC5 | 1.169 | 5.312 | 90.125 |
| PC6 | 0.645 | 2.931 | 93.056 |
| PC7 | 0.455 | 2.068 | 95.124 |
| PC8 | 0.416 | 1.893 | 97.017 |
| PC9 | 0.309 | 1.404 | 98.421 |
| PC10 | 0.130 | 0.589 | 99.010 |
| PC11 | 0.098 | 0.446 | 99.456 |
| PC12 | 0.060 | 0.272 | 99.728 |
| PC13 | 0.041 | 0.185 | 99.912 |
| PC14 | 0.016 | 0.071 | 99.983 |
| PC15 | 0.004 | 0.017 | 100.000 |
| PC16 | 0.000 | 0.000 | 100.000 |
| PC17 | 0.000 | 0.000 | 100.000 |
| PC18 | 0.000 | 0.000 | 100.000 |
| PC19 | 0.000 | 0.000 | 100.000 |
| PC20 | 0.000 | 0.000 | 100.000 |
| PC21 | 0.000 | 0.000 | 100.000 |
| PC22 | 0.000 | 0.000 | 100.000 |

Supplementary Table S 5: Selection differential for mean performance based on the direct and univariate selection on fruit yield

| TRAIT | Xo | Xs | SD | SD (%) | Goal | Success |
|---|---|---|---|---|---|---|
| NNCF | 7.20 | 11.88 | 4.67 | 64.89 | Low | No |
| WNCF | 65.36 | 96.25 | 30.89 | 47.27 | Low | No |
| AWNCF | 8.73 | 7.89 | -0.84 | -9.65 | Low | Yes |
| WUE | 115.84 | 61.66 | -54.18 | -46.77 | Low | Yes |
| NDBF | 57.70 | 54.75 | -2.95 | -5.12 | Low | Yes |
| NDFF | 76.27 | 73.50 | -2.77 | -3.63 | Low | Yes |
| NDBH | 77.80 | 80.88 | 3.08 | 3.96 | Low | No |
| PHYL | 164.45 | 122.82 | -41.63 | -25.31 | Low | Yes |
| TA | 1.42 | 1.31 | -0.11 | -7.54 | Low | Yes |
| NCF | 23.47 | 38.22 | 14.75 | 62.86 | High | Yes |
| TNF | 30.66 | 50.10 | 19.44 | 63.39 | High | Yes |
| WCF | 358.57 | 542.43 | 183.86 | 51.28 | High | Yes |
| TWF | 412.76 | 638.68 | 225.92 | 54.73 | High | Yes |
| AWCF | 14.85 | 14.25 | -0.60 | -4.04 | High | No |
| OAWF | 13.54 | 12.77 | -0.77 | -5.67 | High | No |
| FY | 30957.11 | 47900.84 | 16943.74 | 54.73 | High | Yes |
| TSS | 7.43 | 6.96 | -0.46 | -6.25 | High | No |
| TSS/TA | 5.39 | 5.40 | 0.02 | 0.29 | High | Yes |
| FIRM | 1.67 | 1.54 | -0.13 | -7.86 | High | No |
| L | 54.81 | 54.30 | -0.50 | -0.92 | High | No |
| CHROMA | 44.60 | 44.71 | 0.11 | 0.26 | High | Yes |
| H | 43.07 | 41.83 | -1.24 | -2.88 | High | No |
